# Deep learning and likelihood approaches for viral phylogeography converge on the same answers whether the inference model is right or wrong

**DOI:** 10.1101/2023.02.08.527714

**Authors:** Ammon Thompson, Benjamin Liebeskind, Erik J. Scully, Michael Landis

## Abstract

Analysis of phylogenetic trees has become an essential tool in epidemiology. Likelihood-based methods fit models to phylogenies to draw inferences about the phylodynamics and history of viral transmission. However, these methods are computationally expensive, which limits the complexity and realism of phylodynamic models and makes them ill-suited for informing policy decisions in real-time during rapidly developing outbreaks. Likelihood-free methods using deep learning are pushing the boundaries of inference beyond these constraints. In this paper, we extend, compare and contrast a recently developed deep learning method for likelihood-free inference from trees. We trained multiple deep neural networks using phylogenies from simulated outbreaks that spread among five locations and found they achieve close to the same levels of accuracy as Bayesian inference under the true simulation model. We compared robustness to model misspecification of a trained neural network to that of a Bayesian method. We found that both models had comparable performance, converging on similar biases. We also implemented a method of uncertainty quantification called conformalized quantile regression which we demonstrate has similar patterns of sensitivity to model misspecification as Bayesian highest posterior intervals (HPI) and greatly overlap with HPIs, but have lower precision (more conservative). Finally, we trained and tested a neural network against phylogeographic data from a recent study of the SARS-Cov-2 pandemic in Europe and obtained similar estimates of region-specific epidemiological parameters and the location of the common ancestor in Europe. Along with being as accurate and robust as likelihood-based methods, our trained neural networks are on average over 3 orders of magnitude faster. Our results support the notion that neural networks can be trained with simulated data to accurately mimic the good and bad statistical properties of the likelihood functions of generative phylogenetic models.

## Introduction

Viral phylodynamic models use genomes sampled from infected individuals to infer the evolutionary history of a pathogen and its spread through a population (Holmes and Garnett 1994; Volz et al. 2013). By linking genetic information to epidemiological data, such as the location and time of sampling, these generative models can provide valuable insights into the transmission dynamics of infectious diseases, especially in the early stages of cryptic disease spread when it is more difficult to detect and track (Holmes et al. 1995; Rambaut et al. 2008; Lemey et al. 2009; Pybus et al. 2012; Worobey et al. 2016, 2020; Lemey et al. 2021; Washington et al. 2021; Pekar et al. 2022). This information can be used to inform public health interventions and improve our understanding of the evolution and spread of pathogens. Many phylodynamic models are adapted from state-dependent birth-death (SDBD) processes or, equivalently, state-dependent speciation-extinction (SSE) models (Maddison et al. 2007; FitzJohn 2012; Kühnert et al. 2014; Beaulieu and O’Meara 2016). These birth-death models correspond to the well-known Susceptible-Infectious-Recovered (SIR) model during an exponential growth phase, when nearly all individuals in the population are susceptible to infection (Anderson and May 1979). The simplest SIR models only track the number of susceptible, infected, and recovered individuals across populations over time, with more advanced models also allowing the movement of individuals among localized populations. The phylodynamic models we are interested in track the incomplete transmission tree (phylogeny) of sampled, infected individuals that emerges from host-to-host pathogen spread among populations over space and time. Within this broader context, we will refer to the state as location and the models as location-dependent birth-death (LDBDS) models that include serial sampling of taxa (Kühnert et al. 2016).

Analysts typically fit these birth-death models to data using likelihood-based inference methods, such as maximum likelihood (Maddison et al. 2007; Richter et al. 2020) or Bayesian inference (Kühnert et al. 2016; Scire et al. 2020). Likelihood-based inference relies upon a likelihood function to evaluate the relative probability (likelihood) that a given phylogenetic pattern (i.e., topology, branch lengths, and tip locations) was generated by a phylodynamic process with particular model parameter values. In this sense the likelihood of any possible phylodynamic data set is mathematically encoded into the likelihood as a function of (unknown) data-generating model parameters.

Computing the likelihood requires high-dimensional integration over a large and complex space of evolutionary histories. Analytically integrated likelihood functions, however, are not known for LDBDS models. Methods developers instead use ordinary differential equation (ODE) solvers (Maddison et al. 2007; Kühnert et al. 2016) to numerically approximate the integrated likelihood. These clever approximations perform well statistically, but are too computationally expensive to use with large epidemic-scale data sets. Thus, while Nextstrain (Hadfield et al. 2018) and similar efforts have provided useful visualizations to policy makers during the COVID response, most phylogeographical methods are used forensically, providing insight on the past, and are not used to provide parameter estimates in response to emerging events to inform policy decisions in real-time due to the complexity and long run-times of these models.

As phylodynamic models become more biologically realistic, they will necessarily grow more mathematically complex, and therefore less able to yield likelihood functions that can be approximated using ODE methods. Because of this, phylodynamic model developers tend to explore only models for which a likelihood-based inference strategy is readily available. As a consequence, the lack of scalable inference methods impedes the design, study, and application of richer phylodynamic models of disease transmission, in particular, and richer phylogenetic models of lineage diversification, in general.

To avoid the computational limitations associated with likelihood-based methods, deep learning inference methods that are likelihood-free have emerged as a complementary framework for fitting a wide variety of evolutionary models (Bokma 2006). Deep learning methods rely on training many-layered neural networks to extract information from data patterns. These neural networks can be trained with simulated data as another way to approximate the latent likelihood function (Cranmer et al. 2020). Once trained, neural networks have the benefit of being fast, easy to use, and scalable. Recently, likelihood-free deep learning neural network methods have successfully been applied to phylogenetics (da Fonseca et al. 2020; Suvorov et al. 2020; Nesterenko et al. 2022; Solis-Lemus et al. 2022; Suvorov and Schrider 2022) and phylodynamic inference (Lambert et al. 2022; Voznica et al. 2022).

Here we extend new methods of deep learning from phylogenetic trees (Lambert et al. 2022; Voznica et al. 2022) to explore their potential when applied to phylogeographic problems in geospatial epidemiology. Phylodynamics of birth-death-sampling processes that include migration among locations have been under development for more than a decade (Stadler 2010; Stadler et al. 2012; Kühnert et al. 2014, 2016; Scire et al. 2020; Gao et al. 2022, 2023). Given the added complexity of location-specific dynamics (e.g. location-specific infection rates) and recent successes in deep learning with phylogenetic time trees (Voznica et al. 2022) under state-dependent diversification models (Lambert et al. 2022), we sought to evaluate this approach when applied to viral phylodynamics and phylogeography by including location data when training deep neural networks with phylogenetic trees.

A current limitation of likelihood-free approaches is that it remains unknown how brittle the inference machinery is when the assumptions used for simulation and training are violated (Schmitt et al. 2022). For example, a brittle deep learning method would be more easily misled by model misspecification when compared to a likelihood-based method. Likelihood approaches may have some advantages because the simplifying assumptions are explicit in the likelihood function while for trained neural networks it is difficult to know how those same assumptions implemented in the simulation are encoded in data patterns in the training data and learned network weights. However, with complex likelihood models, there may be unexpected interactions among simplifying assumptions that can result in large biases when applied to real-world data (Gao et al. 2023). Characterizing the relative robustness and brittleness of these two inference paradigms is essential for those who wish to confidently develop and deploy likelihood-free methods of inference from real world data.

To explore relative robustness to model misspecification, we trained multiple deep convolutional neural networks (CNNs) with transmission trees generated from epidemic simulations. We were able to achieve accuracy very close to that of a likelihood-based approach and through several model misspecification experiments show that our CNNs are no more sensitive to model violations than the likelihood approach. Significantly, both methods consistently show similar biases induced by model violations in test data sets. We find that for the models tested here, the migration rate estimates are highly sensitive to misspecification of infection rate and sampling rates, but that estimates of the infection and sampling rates are fairly robust to misspecification of the migration models. We also show that the rate parameter estimates are fairly robust to misspecification of both the number of locations in the model and phylogenetic error. We also estimated prediction intervals for the rate parameters and compared and contrasted their performance to the Bayesian highest posterior density intervals (HPI). We show that they produce intervals that greatly overlap with HPIs in all experiments, but have, on average, wider intervals making them relatively conservative. Finally, we compared a simulation-trained neural network to a recent phylodynamic study of the first wave of the COVID pandemic in Europe (Nadeau et al. 2021) and obtain similar inferences about the dynamics and history of SARS-CoV-2 in the European clade.

## Methods

First, we define the SIR model we assume here that is approximately equivalent to the LDBDS model (Kühnert et al. 2016). Following that, is a description of the simulation method to generate the training, validation, and test data sets of phylogenies under the model. The simulation and data processing pipeline is shown in Figure 1. We next describe our implementation of simulation-trained deep learning inference with convolutional neural networks (CNN) as well as a likelihood-based method using Bayesian inference. We then describe our methods for measuring and comparing their performance when tested against data sets generated by simulations under the inference model as well as several data sets simulated under models that violate assumptions of the inference model. Finally, we describe how we tested our simulation-trained CNN against a real-world data set.

**Figure 1:**
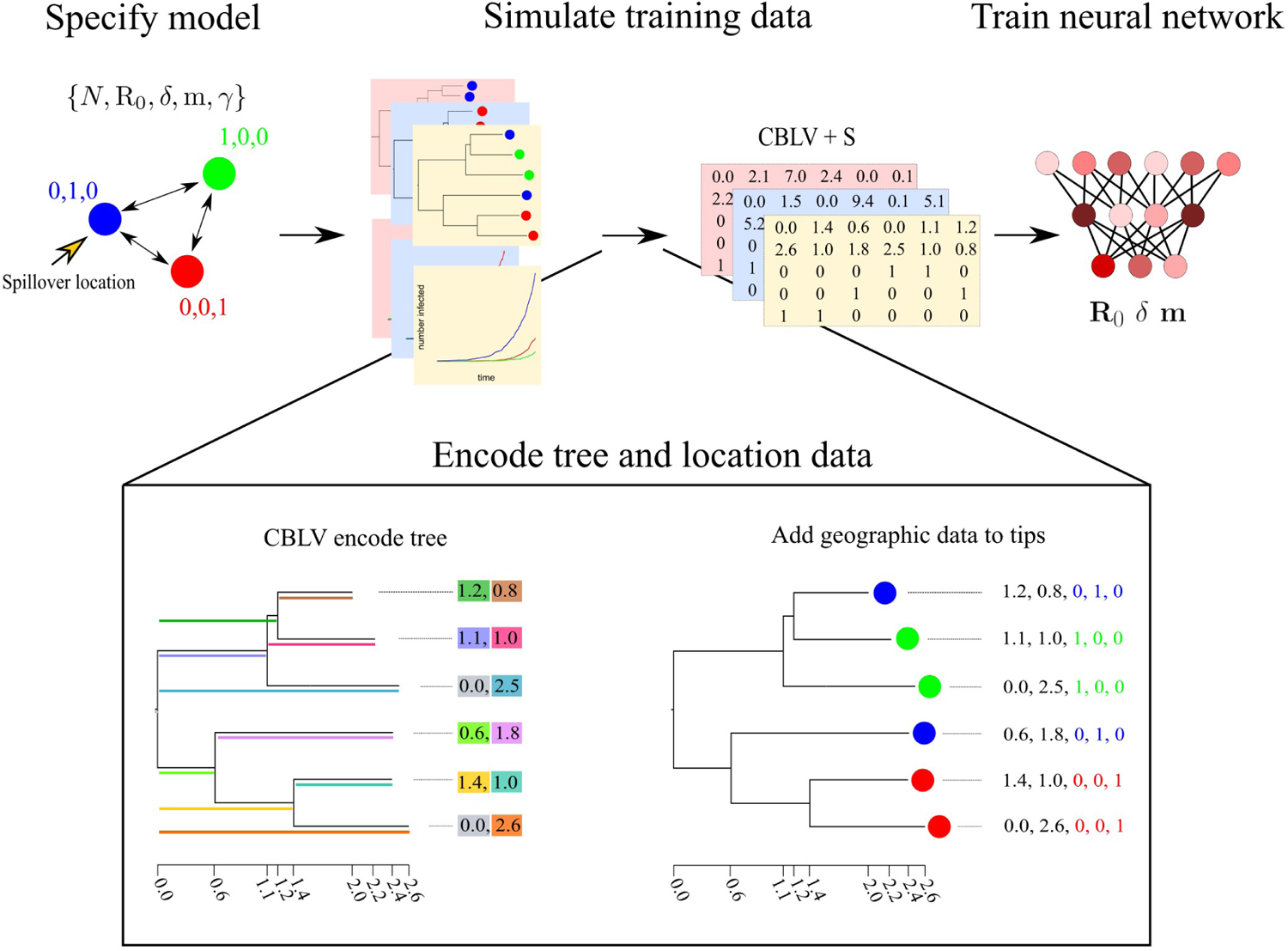
Simulation and tree encoding pipeline for generating training data. 1) Specify a model, for example an SIR model with serial sampling and migration among three locations (colored circles). 2) Run simulations of outbreaks under the model to generate population trajectories and phylogenetic trees. 3) Encode trees and location data into the Compact Bijective Ladderized Vector + States (CBLV+S) format. 4) Train the neural network with CBLV+S training data.

### Model definition

We first define a general location-dependent SIR stochastic process used for simulations and likelihood function derivation in the format of reaction equations we specified in MASTER (Vaughan and Drummond 2013). Reaction equations 1 through 4 specify the SIR compartment model with migration and serial sampling where *S*, *I*, and *R* denote the number of individuals in each compartment. The *S* and *I* compartments are indexed by geographic location using *i* and *j*. *N_i_*is the total population size in location *i* and *N_i_* = *S_i_* + *I_i_* + *R_i_*. To simplify notation, we consider all local recoveries to lead to the same global compartment and absorbing state, *R*. The symbols for each rate parameter is placed above each reaction arrow.

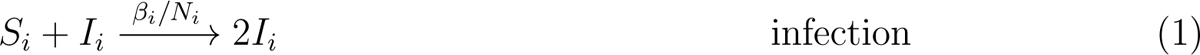

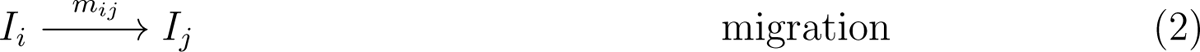

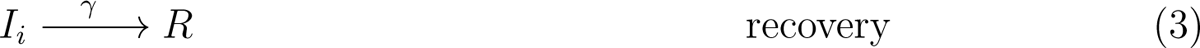

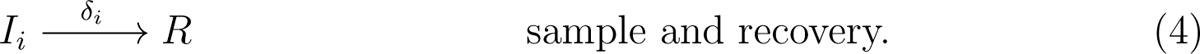

We parameterize the model with the basic reproduction number in location *i*, *R*_0_*_i_*, which is related to *β_i_* and *δ_i_* by equation 5,

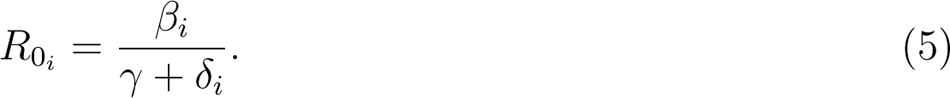

In particular, our study considers a location-independent SIR (LISIR) model with sampling that assumes *R*_0_*_i_* was equal among all locations, and a location-dependent (LDSIR) model with sampling that assumes *R*_0_*_i_* varied among locations. During the exponential growth phase of an outbreak, the LISIR and LDSIR models are equivalent to the location-independent birth-death-sampling (LIBDS) and location-dependent birth-death-sampling (LDBDS) models, respectively, that are often used in viral phylogeography (Kühnert et al. 2014, 2016; Douglas et al. 2021).

Each infectious individual transitions to recovered at rate *γ*. We assumed that sampling a virus in an individual occurs at rate *δ_i_* in location *i* and immediately removes that individual from the infectious compartment and places them in the recovered compartment. Thus the effective recovery rate in location *i* is *γ* + *δ_i_*. The above reactions correspond to the following coupled ordinary differential equations.

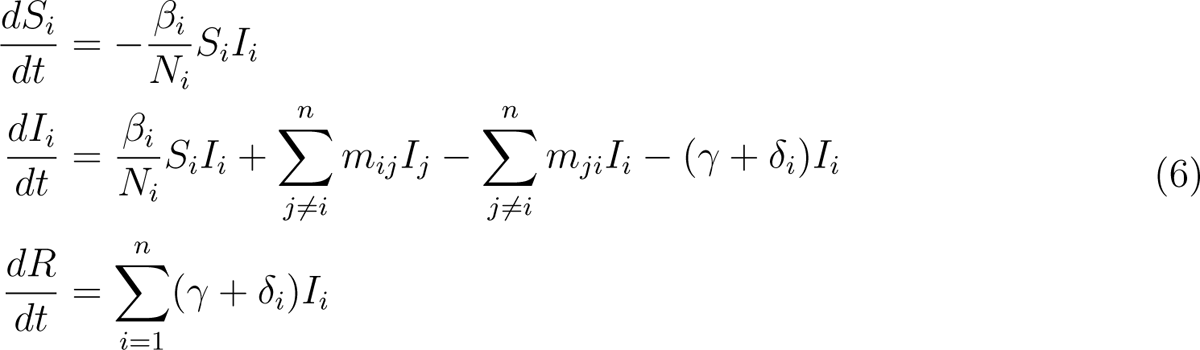

When the migration rate is constant among locations and the model is a location-independent SIR model, or equivalently, LIBDS, and we set *S_i_*(*t* = 0) *≈ N_i_* at the beginning of the outbreak, the equation set 6 reduces to

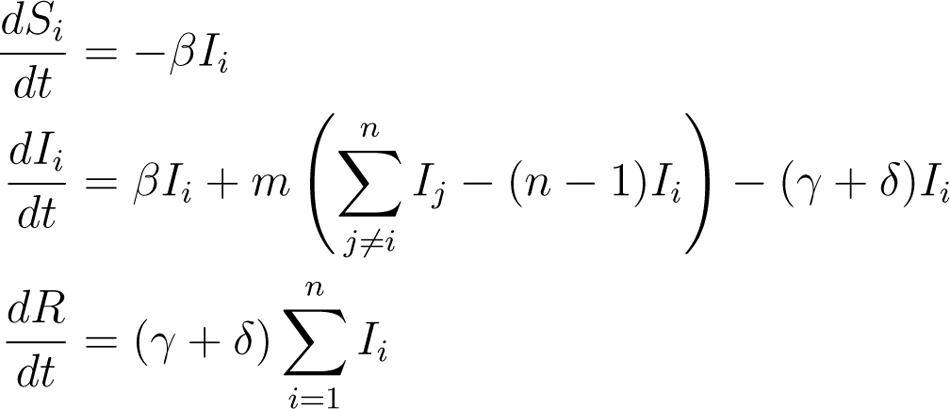

The number of infections and the migration of susceptible individuals is at negligible levels on the timescales investigated here. The infection rate is, therefore, approximately constant and the migration of susceptible individuals can be safely ignored requiring only migration of infectious individuals to be simulated.

At the beginning of an outbreak, it is often easier to know the recovery period from clinical data than the sampling rate which requires knowing the prevalence of the disease. Therefore, we treat the average recovery period as a known quantity and use it to make the other two parameters (the sampling rate and the basic reproduction number R_0_) identifiable. This was done by fixing the corresponding rate parameter in the likelihood function to the true simulated value for each tree, and by adding the true simulated value to the training data for training the neural network.

### Simulated training and validation data sets

Epidemic simulations of the SIR+migration model that approximates the LIBDS process were performed using the MASTER package v. 6.1.2 (Vaughan et al. 2014) in BEAST 2 v. 2.6.6 (Bouckaert et al. 2019). MASTER allows users to simulate phylodynamic data sets under user-specified epidemiological scenarios, for which MASTER simultaneously simulates the evolution of compartment (population type) sizes and tracks the branching lineages (transmission trees in the case of viruses) from which it samples over time. We trained neural networks with these simulated data to learn about latent populations from the shape of sampled and subsampled phylogenies. In addition to the serial sampling process, at the end of the simulation 1% of infected lineages were sampled. In MASTER this was approximated by setting a very high sampling rate and very short sampling time such that the expected number sampled was approximately 1%. This final sampling event was required to make a 1-to-1 comparison of the likelihood function used for this study (see Likelihood method description below) which assumes at least one extant individual was sampled to end the process. Coverage statistics from our MCMC samples closely match expectations (see Likelihood method description below; SI Figure 2 C). Simulation parameters under LIBDS and LDBDS models for training the neural network under the phylogeography model were drawn from the following distributions:

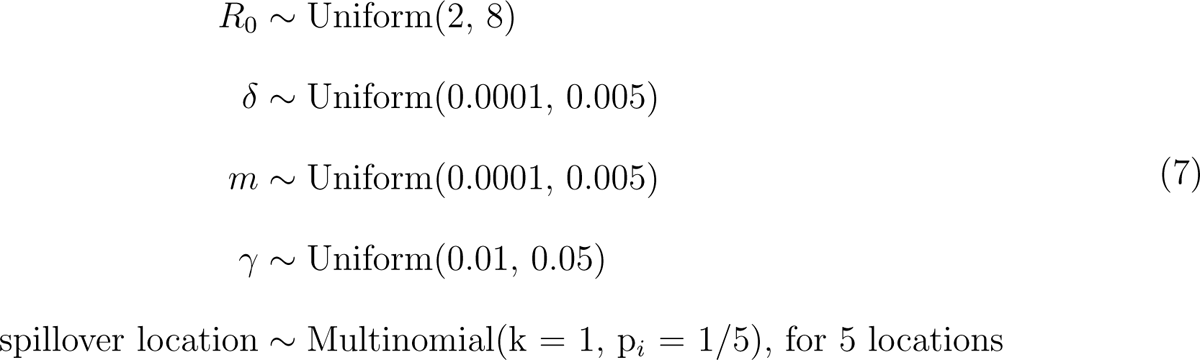

**Figure 2:**
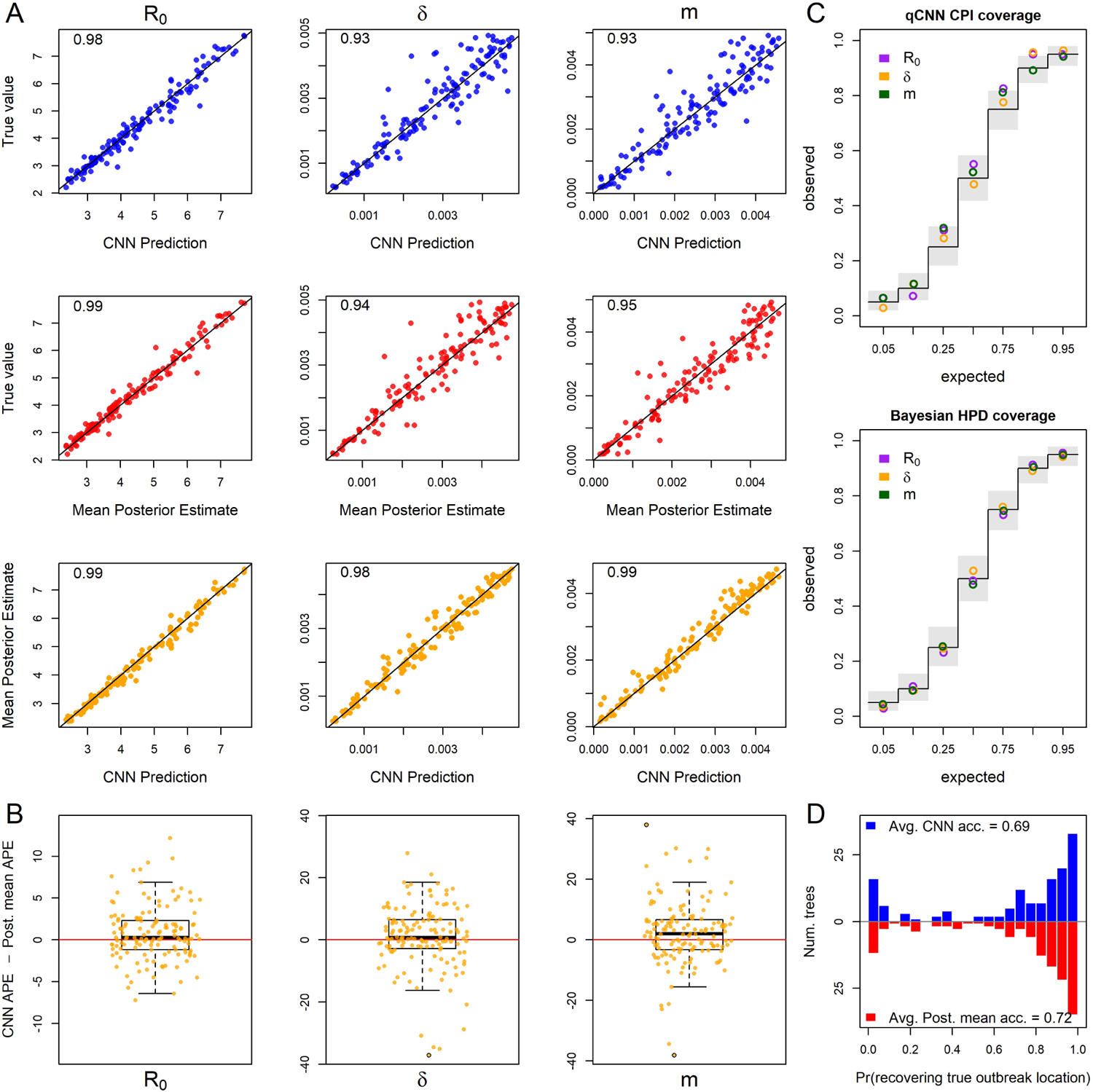
Inference under the true simulating model. (A) Scatterplot of CNN predictions and posterior mean estimates from Bayesian analyses against the true values (top two rows in blue and red respectively) of the basic reproduction number, R_0_, the sampling rate, *δ*, and the migration rate, m for 138 test trees. In the upper-left corners of the scatter plots are the correlations of the plotted data. The bottom row in orange shows scatter plots of the CNN estimates against the posterior mean estimates for the same trees. (B) The difference in the absolute percent error (APE) of estimates for the two inference methods. Boxes show the inner 50% quantile of the data while whiskers extend 1.5 IQR. Dots with black circles were truncated to 2*×* the length of whiskers for visualization purposes. (C) Coverage plots show the expected frequency of coverage for each of the categories and the observed frequencies (black steps and colored circle respectively). Gray boxes are the expected 95% confidence intervals at each of the expected coverage values which follows a Beta((*n*+1)*q, n−*(*n*+1)*q*+1) distribution. (D) Histograms of the probabilities of inferring the correct outbreak origin location.

All five locations had initial population sizes of 1,000,000 susceptible individuals and one infected individual in a randomly sampled spillover location. Simulations were run for 100 time units or until 50,000 individuals had been infected to restrict simulations to the approximate exponential phase of the outbreak. For the experiments comparing the CNN to the likelihood-based method under the LIBDS model, if this population threshold was reached the simulation was rejected. This criterion was not enforced for simulations under the LDBDS model. This ensured the LIBDS model used in the likelihood-based analyses are equivalent to more complex density-dependent SIR models. After simulation, trees with 500 or more tips were uniformly and randomly downsampled to 499 tips and the sampling proportion was recorded for training the neural networks and to adjust estimates of *δ*.

We simulated 410,000 outbreaks under these LIBDS settings to generate the training, validation, and test sets for deep learning. Any simulation that generated a tree with less than 20 tips was discarded, leaving a total of 111,157 simulated epidemiological data sets. Of these, 104,157 data sets were used to train and 7,000 were used to validate and test each CNN. A total of 193,110 LDBDS data sets were simulated, with 186,110 used to train and 7,000 used to validate and test the LDBDS CNNs.

To make phylodynamic inferences about the first wave of the SARS-CoV-2 epidemic in Europe we used the LDBDS model on the data set from Nadeau et al. (2021). Training simulation parameters for the LDBDS process were drawn from the same distributions as LIBDS except R_0_ which was unique for each location. We assume that the variability of R_0_ among different pathogens (simulated outbreaks) is greater than the variability of the same pathogen’s R_0_ among different locations within the same simulation. To implement this assumption, all R_0_ was drawn from a joint distribution to narrow the magnitude of differences among locations within simulations to be within 6 of each other but expand the magnitude of differences between simulations to range from 0.9 to 15:

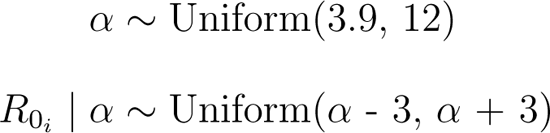

For the empirical analysis, population sizes at each location were also set to 500,000 and instead of running the simulations for 100 time units, time was scaled by the recovery period, 1*/γ*, and was drawn from a uniform distribution:

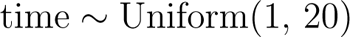

### Simulated test data sets with and without model misspecification

All simulation models used for training and testing are listed in Table 1. We first simulated a test set of 138 trees under the training model to compare the accuracy of the CNN and the likelihood-based estimates when the true model is specified. These data sets were simulated by random draws of parameter values from the same distributions described above for generating the training data set.

**Table 1:**
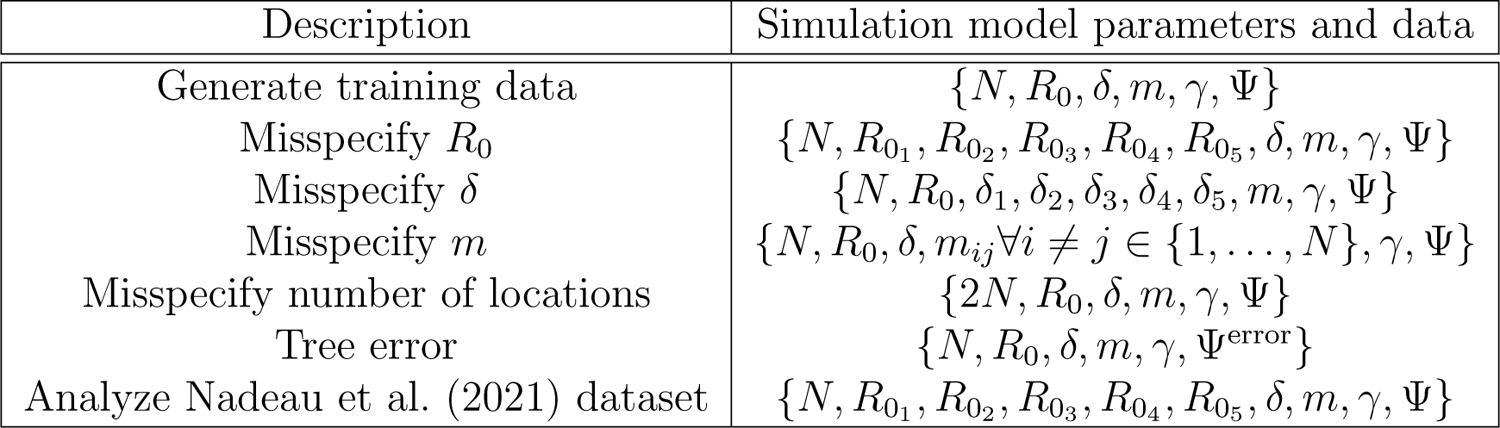
Models used in this study. All simulations assume an SIR compartmental epidemic model. N = 5 is the number of locations, R_0_ is the basic reproduction number, *δ* is the sampling rate, m is the migration rate, *γ* is the recovery rate (treated as data), and Ψ is the phylogenetic tree + locations (also treated as data).

Sensitivity to model misspecfication for each of the three rate parameters, *R*_0_, *δ*, and *m*, was tested. All sensitivity experiments used the same LIBDS model for inference for both the CNN and the Likelihood-based methods. Sensitivity experiments were conducted by simulating a test data set of trees that were generated by an epidemic process that was more complex than or different from the LIBDS model.

The tree data set for the misspecified R_0_ experiment consisted of simulating outbreaks where each location had a unique R_0_ drawn from the same distribution as above. Likewise, the misspecified sampling model test set was generated by simulating outbreaks where each location had a unique sampling rate, *δ*, drawn from the same distribution used for the global sampling rate described above. For the misspecified migration model, a random pair of coordinates, each drawn from a uniform(0,5) distribution in a plane, were generated for the five locations, and a pairwise migration rate was computed such that pairwise migration rates were symmetric and proportional to the inverse of their euclidean distances and the average pairwise migration rate was equal to a random scalar which was also drawn from a uniform distribution (see equations 7 above).

The tree set for the misspecified number of locations experiment was generated by simulating outbreaks among ten locations instead of five. After simulations, six locations were chosen at random and re-coded as being sampled from the same location.

To generate a test set where the time tree used for inference has incorrect topology and branch lengths, we implemented a basic pipeline of tree inference from simulated genetic data to mimic a worst case real world scenario. We simulated trees under the same settings as before. Phylogenetic error was introduced in two ways: the amount of site data (short sequences) and misspecification of the DNA sequence evolution inference model using seq-gen V. 1.3.2 (Rambaut and Grassly 1997). We simulated the evolution of a 200 base-pair sequence under an HKY model with *κ* = 2, equal base frequencies and 4 discretized-gamma(2, 2) rate categories for among site rate variation. The simulated alignment as well as the true tip dates (sampling times) was then used to infer test trees. Test tree inference was done using IQ-Tree v. 2.0.6 (Minh et al. 2020) assuming a Jukes-Cantor model of evolution where all transition rates are equal. The inference model also assumed no among-site rate variation. The number of shared branches between the true transmission tree and the test tree inferred by IQ-Tree was measured using gotree v. 0.4.2 (Lemoine and Gascuel 2021). Polytomies were resolved using phytools (Revell 2012) and a small, random number was added to each resolved branch. These trees were then used for likelihood inference and CNN prediction.

### Deep learning inference method

The resulting trees and location metadata generated by our pipeline were converted to a modified CBLV format (Voznica et al. 2022), which we refer to as the CBLV+S (+State of character, *e.g.* location) format (Figure 1). The CBLV format uses an in-order tree traversal to translate the topology and branch lengths of the tree into an 2 x n matrix where n is the maximum number of tips allowed for trees. The matrix is initialized with zeroes. We then fill the matrix starting with the root then proceed to the tip with largest root-to-tip distance rather than starting with that tip as in Voznica et al. (2022). We chose this to separate the the zero value of the root age from the zeroes used to pad matrices where the tree has less than the maximum number of tips, though we expect this to make marginal to no difference in performance. The CBLV representation gives each sampled tip a pair of coordinates in ‘tree-traversal space’. Our CBLV+S format associates geographic information corresponding with each sampled taxon by appending each vector column with a one-hot encoding vector of length *g* states to yield a (2 + *g*) *× n* CBLV+S matrix. The CBLV+S format allows for multiple characters and/or states to be encoded, extending the single binary character encoding format introduced by Lambert et al. (2022). Our study uses CBLV+S to encode a single character with *g* = 5 location-states. In addition to the the CBLV+S data, we also include a few tree summary statistics and known simulating parameters; the number of tips, mean branch length, the tree height and the recovery rate and the subsampling proportion. Trees were rescaled such that their mean branch length was the default for phylodeep (Voznica et al. 2022) before training and testing of the CNN. The mean pre-scaling branch length and tree heights were also fed into the neural networks. Trees were not rescaled for the likelihood-based analysis. Recall that tree height did not vary for the LIBDS CNN training set but did for the LDBDS training set (see simulation time settings above). Varying the time-scale for the LDBDS model was necessary for analyzing real world data where time-scales of outbreaks can vary considerably.

Our CNNs were implemented in Python 3.8.10 using keras v. 2.6.0 and tensorflow-gpu v. 2.6.0. (Chollet; Abadi et al. 2016). CNNs consist of one or more layers specifically intended for structural feature extraction. CNNs utilize a filter, akin to a sliding window, that executes a mathematical operation (convolution) on the input data. When dealing with structured data like the CBLV+S matrix, multiple 1D filters slide across the matrix’s columns, embedding each scanned window into an N-dimensional vector representation. This architectural design imparts CNNs with translation invariance, enabling them to recognize and learn repeating patterns throughout the input space, regardless of their specific location. Stacking multiple convolutional layers enables CNNs to decipher hierarchical structures within the data. See Alzubaidi et al. (2021) and Khan et al. (2020) for reviews of the subject.

For each model, LIBDS and LDBDS, we designed and trained two CNN architectures, one to predict epidemiological rate parameters and the other to predict the outbreak location resulting in four total CNNs trained by two training data sets (LIBDS and LDBDS). We used the mean-squared-error for the regression neural loss function in the network trained to estimate epidemiological rates, and the categorical cross-entropy loss function for the categorical network trained to estimate outbreak location. We assessed the performance of the network by randomly selecting 5,000 samples for validation before each round of training. We measured the mean absolute error and accuracy using the validation sets. We used these measures to compare architectures and determine early stopping times to avoid overfitting the model to the training data. We also added more simulations to the training set until we could no longer detect an improvement in error statistics. After comparing the performance of several networks, we found that the CNN described in SI Figure S1 performed the best. In brief, the networks have three parallel sets of sequential convolutional layers for the CBLV+S tensor and a parallel dense layer for the priors and tree statistics. The three sets of convolution layers differed by dilation rate and stride lengths. These three segments and the dense layer were concatenated and then fed into a segment consisting of a sequential set of dense layers, each layer gradually narrowing to the output size to either three or five for the rates and origin location networks, respectively, for the LIBDS model, and seven and five for the seven rates and five locations, respectively, for the LDBDS model.

All layers of the CNN used rectified linear unit (ReLU) activation functions. We used the Adam optimizer algorithm for batch stochastic gradient descent (Kingma and Ba 2017) with batch size of 128. We selected the number of epochs by monitoring the mean absolute error and accuracy of the validation data set. This set was not used in training or testing. These metrics suggested stopping after 15 epochs for the regression network and ten epochs for the root location network would maximize accuracy/minimize error for out-of-sample test data. The output layer activation for the network that predicted the R_0_*, δ* and m parameters was linear with three nodes. For the output layer predicting the outbreak location the activation function was softmax with five nodes for the five locations. The input layer and all intermediate (latent) layers were the same for all four networks, namely the CBLV+S tensor and the recovery rate, mean branch lengths, tree height and number of tips in the tree. The LDBDS neural network was trained with simulated trees where R_0_*_i_* varied among locations and had an output layer with seven nodes; five for the each location’s R_0_*_i_* and a node each for the sampling rate and the migration rate. We tested networks with max-pooling layers between convolution layers as well as dropout at several rates and found no improvement or a decrease in performance.

### Likelihood-based method of inference

We compared the performance of our trained phylodynamic CNN to likelihood-based Bayesian phylodynamic inferences. We specified LIBDS and LDBDS Bayesian models that were identical to the LIBDS and LDBDS simulation models that we used to train our CNNs. The most general phylodynamic model in the birth-death family applied to epidemiolgoical data is the state-dependent birth-death-sampling process (SDBDS; (Kühnert et al. 2016; Scire et al. 2020)), where the state or type on which birth, death, and sampling parameters are dependent is the location in this context. The basic model used for experiments here is a phylogeographic model that is similar to the serially sampled birth-death process (Stadler 2010) where rates do not depend on location, which we refer to as the LIBDS model. The death rate, *µ*, is equivalent to the recovery rate, *γ*, in SIR models. Standard phylogenetic birth-death models assume the birth and death rates, *λ* and *µ*, are constant or time-homogeneous, while the SIR model’s infection rate is proportional to *β* and *S* and varies with time as *S* changes. However, when the number of infected is small relative to susceptible people, as in the initial stages of an outbreak, the infection rate, *β*, is approximately constant and approximately equal to the birth rate *λ*;

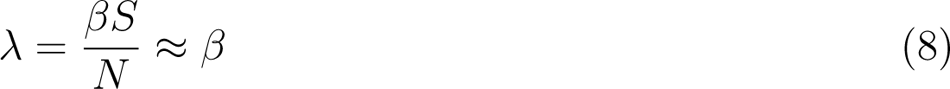

The joint prior distribution was set to the same model parameter distributions that were used to simulate the training and test sets of phylogenetic trees in the first section with *γ* treated as known and the proportion of extant lineages sampled, *ρ*, set to 0.01 as in the simulations. The likelihood was conditioned on the tree having extant samples (*i.e.* the simulation ran for the allotted time without being rejected). All simulated trees in this study had a stem branch and the outbreak origins were inferred for the parent node of the stem branch.

We used Markov chain Monte Carlo (MCMC) to simulate random sampling from the posterior distribution implemented in the TensorPhylo plugin (https://bitbucket.org/mrmay/tensorphylo/src/master/) in RevBayes (Höohna et al. 2016). After a burnin phase, a single chain was run for 7,500 cycles with 4 proposals per cycle and at least 100 effective sample size (ESS) for all parameters. If the effective sample size (ESS) was less than 100, the MCMC was rerun with a higher number of cycles. We also analyzed the coverage of the 5, 10, 25, 50, 75, 90, and 95% HPI to verify that our simulation model and inference model are the same and that the MCMC simulated draws from the true posterior distribution. Bayesian phylogeographic analysis recovered the true simulating parameters at the expected frequencies (Figure 2 C), thus validating the simulations were working as expected and confirming that the MCMC was accurately simulating draws from the true posterior distribution.

### Quantifying errors and error differences

We measure the absolute percent error (APE) of the predictions from the CNN and the mean posterior estimate (MPE) of the likelihood-based method. The formula for APE of a prediction/estimate, *y*^estimate^, of *y*^truth^ is

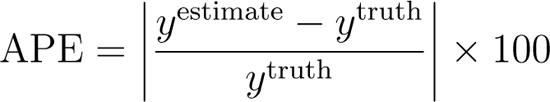

The Bayesian alternative to significance testing is to analyze the posterior distribution of parameter value differences between groups. In this framework, the probability that a difference is greater than zero can be easily interpreted. We therefore used Bayesian statistics to infer the median difference in error between the CNN and likelihood-based methods and the increase in median error of each method when analyzing misspecified data compared to when analyzing data simulated under the true inference model.

We used Bayesian inference to quantify population error by performing three sets of analyses: (1) inferred the population median APE under the true model (this will be the reference group for analysis 3), (2) the effect of inference method — CNN or likelihood-based (Bayesian) — on error by inferring the median difference between the CNN estimate and the likelihood-based estimate, (3) the effect of misspecification on error for each parameter by comparing the median error of estimates under misspecified experiments and the reference group defined by analysis 1. See SI Figures S3 - S13 and SI Table S1 for summaries and figures for all analyses for this section.

To infer these differences between groups we used the R package BEST (Meredith and Kruschke). BEST assumes the data follow a t-distribution parameterized by a location parameter, *µ*, a scale parameter, *σ*, and a shape parameter, *ν*, which they call the “normality parameter” (*i.e.* if *ν* is large the distribution is more Normal). Because the posterior distribution does not have a closed form, BEST uses Gibbs sampling to simulate draws from the posterior distribution. 20,000 samples were drawn from the posterior distribution for each BEST analysis. BEST uses automatic posterior predictive checks to indicate that a model adequately describes the data distributions. Posterior predictive checks indicate the BEST model adequately fits each data set analyzed below.

### Inferring the median APE

Before inferring differences between groups, we inferred the population median APE for predictions of R_0_, *δ*, and *m* from test data simulated under the inference model using the CNN and likelihood-based methods. Histograms of the sampled log-transformed APE appears to be symmetric with heavy tails so we fit the log APE to the BEST model. This implies that the sampled APE scores are drawn from a log-t distribution. The log-t distribution has a mean of *∞* and median of *e^µ^*, we therefore focus our inference on estimating posterior intervals for the population median APE from the sampled APE values for each parameter estimated by the CNN method and likelihood-based method which we denote APE^CNN^, and APE^Like^ respectively. The data analyzed here and likelihood assumed by BEST is

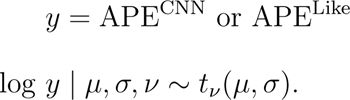

The priors were set to the vague priors that BEST provides by default,

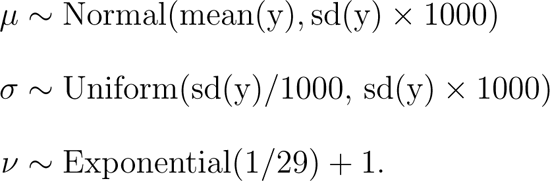

95% HPI for the median APE, *µ*°, was estimated by the following transformation of simulated draws from the posterior distribution

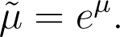

In summary, the results we present are 95% HPI from the posterior distributions of the median error, *µ*°.

### Inferring the relative accuracy of the CNN and likelihood-based method

To quantify the difference in error between the CNN and the likelihood-based method, we fit the difference in sampled APE scores, ΔAPE, between the CNN method and the likelihood-based method to the BEST model. Histograms of ΔAPE appear symmetric with weak to strong outliers making the BEST model a good candidate for inference from this data. The data and likelihood are

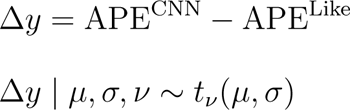

We used the same default priors as above.

Because, Δ*y* is not log-transformed, it is drawn from a t-distribution and the marginal posterior of the parameter *µ* is an estimate of the population mean, *µ^d^*. Because the mean and the median are equivalent for a t-distribution, we again report the posterior distribution of the median difference, *µ*°*^d^* to simplify the results.

In summary, the results we present are 95% HPI from the posterior distribution of the median difference between the two methods, *µ*°*^d^*.

When comparing CNN to the likelihood-based approach, positive values for *µ*°*^d^* indicate the CNN is less accurate, and negative indicate the likelihood-based estimates less accurate. We emphasise that this quantity is the median difference in contrast to the difference in medians, Δ*µ*°, reported in the next section.

### Inferring sensitivity to model misspecification

Finally, to quantify the overall sensitivity of each rate parameter to model misspecification under each inference method, we infer the difference in median APE, *µ*° of predictions under a misspecified model relative to predictions under the true model. In other words we are inferring differences in medians between experiments. For example, to infer the sensitivity of the CNN’s inference of the sampling rate, *δ*, to phylogenetic error, we inferred the difference between the median APE of the CNN’s predictions for misspecified trees and the median APE of CNN predictions for true trees. The data is concatenated as below.

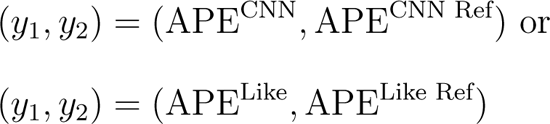

We inferred the difference between group median APE scores, denoted Δ*µ*°, by assuming that the model parameters conditioned on the observed APE from the two groups, *y*_1_ and *y*_2_, follow a posterior distribution that is proportional to

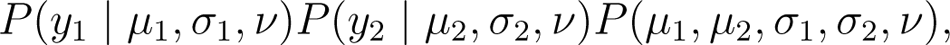

where log *y*_1_ and log *y*_2_ follow t distributions with means *µ*_1_ and *µ*_2_ and standard deviations *σ*_1_ and *σ*_2_, respectively while sharing a common normality parameter, *ν*.

The posterior sample of Δ*µ*° is obtained by transforming samples from the joint marginal posterior distribution of *µ*_1_ and *µ*_2_ with the following equation,

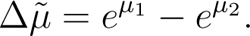

The two components of the likelihood are each t-distributed and share the *ν* parameter which means we assume both samples are drawn from a similarly shaped distribution (similarly heavy tails).

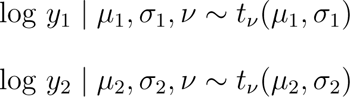

The prior distribution for the parameters of the model were set to the defaults for BEST,

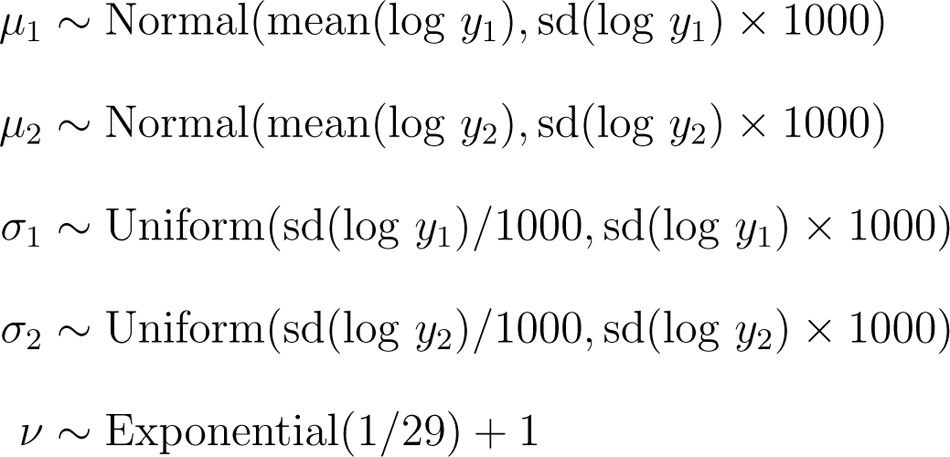

As before, interpretation of the posterior distribution of the difference in medians is straightforward: the more positive the difference in median APE from the misspecified model test set and the median APE from the true model test set, the more sensitive the parameter is to model misspecification in the experiment.

### CNN uncertainty quantification

We used conformalized quantile regression (CQR) to construct calibrated probability intervals (CPI), ensuring accurate predictive coverage (Lei et al. 2018; Romano et al. 2019; Sousa et al. 2022; Vovk et al. 2022; Angelopoulos et al. 2023). CQR is implemented in two stages: first a network is trained to predict conditional quantiles, then a hold-out simulated dataset is used to estimate bias adjustment terms to ensure correct coverage on future data *i.e.* 95% intervals contain the true value 95% of the time for test data.

To implement quantile regression with a neural network and predict lower and upper quantiles, we adjusted the general network architecture used for point estimates above to have two outputs each with a mean pinball loss function instead of the mean squared error,

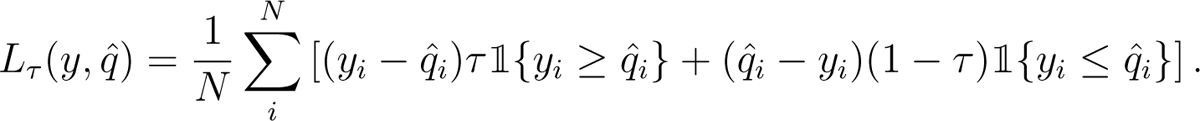

Here, *y* is the label or true parameter value (not a quantile) and *q*^ is the trained neural network’s prediction of a given quantile. *τ* is the quantile level and is equal to 1 *− α*, where *α* is the mis-coverage rate, or the probability the true value is not below the quantile. To estimate inner quantiles with miscoverage rate *α*, the lower quantile output was set to predict the *α/*2 quantile for each rate parameter and the other layer to predict the 1 *− α/*2 upper quantile (Steinwart and Christmann 2011) (SI Figure S2). We refer to CNNs of this type as qCNN. Though often close, these inner quantiles are not guaranteed to have the correct coverage on test data sets (Figure 3) necessitating the calibration (conformalization) step (Romano et al. 2019).

**Figure 3:**
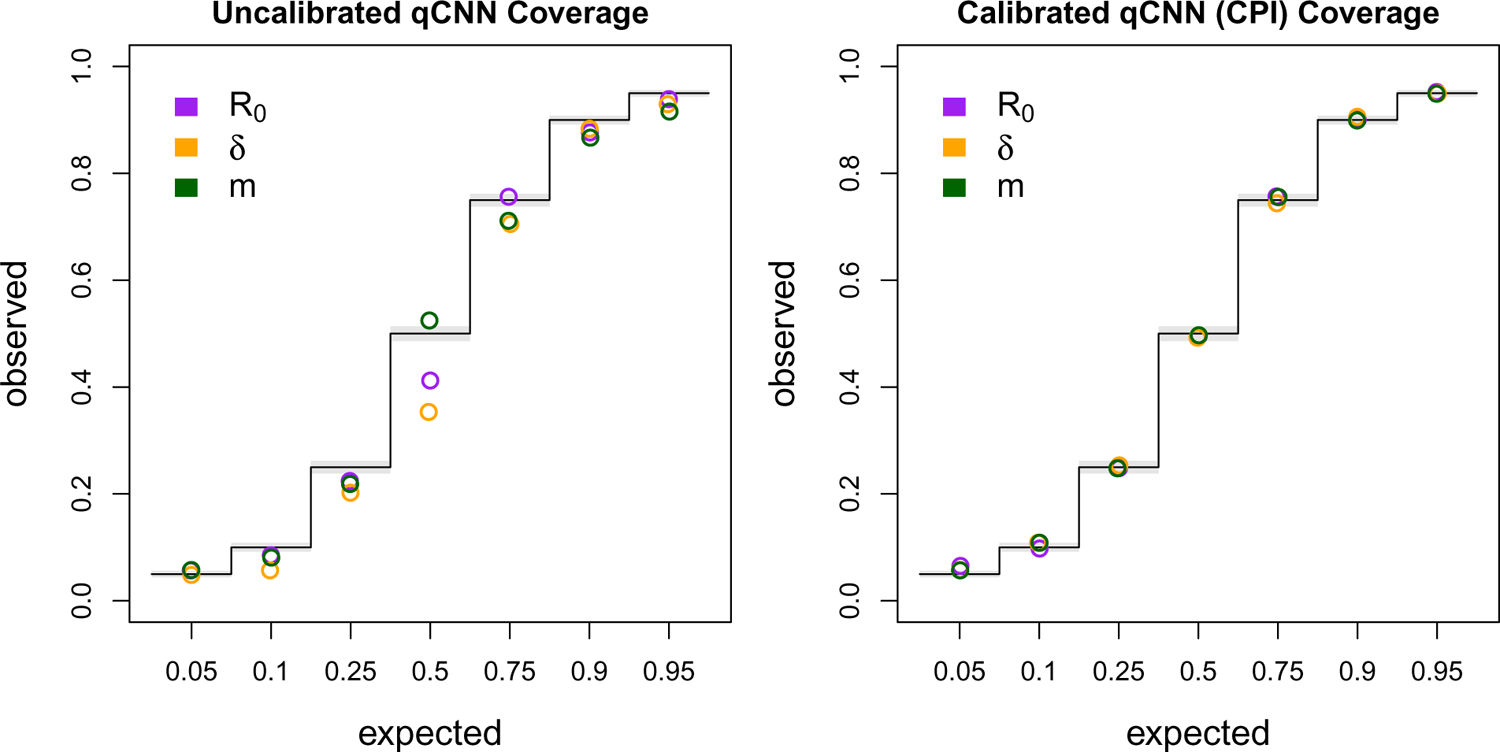
Coverage of uncalibrated qCNN quantile predictions (left) and calibrated qCNN which produce “calibrated probability intervals” (CPI) on the right. The observed coverage of 5,000 samples tested at seven different predicted coverage levels (labeled horizontal). See Figure 2 C for more details on coverage plots.

To calibrate the predictions of quantile regression neural networks, CQR finds an adjustment term for each quantile through computing a non-comformity score, such as the distance of the predicted value from the predicted quantile. If the estimated quantile is well calibrated, then the same quantile of the scores in a calibration set will be zero. If the estimated quantile is, for example, too high then too high a proportion of the labels will fall below the estimated quantile and the empirical quantile, *Q*, of the nonconformity score *y − q*^ at 1 *− α/*2 will be negative. In other words it will over cover the calibration set. *Q* thus becomes the adjustment term for calibrating the qCNN’s quantile estimate (equations 9, and 10) by simply adding the term to the corresponding estimated quantile as shown in equation 11.

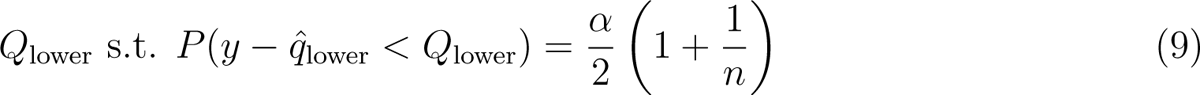

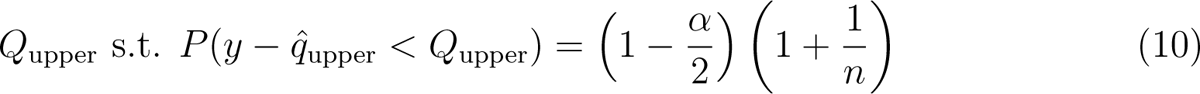

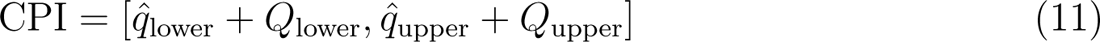

Note that the quantiles of the score for finite sample sizes require adjustment by (1 + <u>1/n</u>) where *n* is the number of samples in the calibration set (Romano et al. 2019).

We simulated 108,559 more datasets (trees) to estimate the calibration amounts for the upper and lower qCNN-estimated quantiles. After calibration through conformalization, we clipped intervals to the prior boundary for intervals that extended beyond the prior distribution’s range. To examine the consistency of quantile regression for neural networks trained on different quantiles we trained seven different quantile networks to predict the same quantiles used for validating our Bayesian analysis and simulation model: *{*0.05, 0.25, 0.5, 0.75, 0.9, 0.95*}*. We checked the coverage of these adjusted CPIs on another simulated test dataset of 5,000 trees.

### Real data

We compared the inferences of a LDBDS simulation trained neural network to that of a phylodynamic study of the first COVID wave in Europe (Nadeau et al. 2021). These authors analyzed a phylogenetic tree of viruses sampled in Europe and Hubei, China using a location-dependent birth-death-sampling model in a Bayesian framework using priors informed by myriad other sources of information. We simulated a new training set of trees under an LDBDS model where R_0_*_i_* depends on the geographic location, and the sampling process only consists of serial sampling and no sampling of extant infected individuals. We estimated 95% CPIs for model parameters with a simulated calibration dataset of 101,219 trees using CQR as above and confirmed accurate coverages with another dataset of 5,000 trees.

We then analyzed the whole tree from Fig. 1 in (Nadeau et al. 2021) as well as the European clade which Nadeau et al. (2021) labeled as A2 in the same figure. We note that our simulating model is not identical to the inference model used in (Nadeau et al. 2021). We model migration with a single parameter with symmetrical migration rates among locations and all locations having the same sampling rate. Nadeau and colleagues parameterize the migration process with asymmetric pairwise migration rates and assume location-specific sampling rates. We also do not include the information the authors used to inform their priors as that requires an extra level of simulation and training on top of simulations done here, and is thus beyond the scope of this study.

The time tree from (Nadeau et al. 2021) was downloaded from GitHub (https://github.com/SarahNadeau/cov-europe-bdmm). The recovery rate assumed in (Nadeau et al. 2021) was 0.1 days*^−^*^1^ which was set to 0.05 to bring the recovery rate to within the range of simulating values used to train the CNN. Consequently, the branch lengths of the tree were then scaled by 2. The number of tips, tree height, and average branch lengths were measured from the rescaled trees and fed into the network. The full tree and A2 clade were then analyzed using the LDBD CNN and compared to the posterior distributions from (Nadeau et al. 2021).

### Hardware used

Simulations were run on a 16 core Intel(R) Xeon(R) Platinum 8175M CPU @ 2.50GHz. For each simulation, an XML file with random parameter settings was generated using custom scripts. These XML files were the inputs for MASTER which was run in the BEAST2 platform. Neural network training and testing and predictions were conducted on an 8 core Intel(R) Core(TM) i7-7820HQ CPU @ 2.90GHz laptop with a NVIDIA Quadro M1200 GPU for training.

## Results

### Comparing deep learning to likelihood

Our first goal in this study was to train a CNN that produced phylodynamic parameter point estimates that were as accurate as likelihood-based Bayesian posterior mean estimates under the true model. This will serve as a reference for quantifying level of sensitivity in our misspecification experiments. Using viral phylogenies like those typically estimated from serially sampled DNA sequences, we focused on estimating important epidemiological parameters – the reproduction number, *R*_0_, the sampling rate, *δ*, the migration rate, *m*, and the outbreak origin.

Our CNN produced estimates that are as accurate as the mean posterior estimates (MPE) under the true simulating model. We compared the absolute percent error (APE) of the network predictions to the APE of the MPE of the Bayesian location-independent birth-death-sampling (LIBDS) model (Figure 2). The APE is straight-forward to interpret, e.g. an APE of *<* 10 means the estimate is within 10 percentage points (ppts) of the true value. For the three epidemiological rate parameters, *R*_0_, *δ* and *m*, both methods made very similar predictions for the 100 time tree test set (Figure 2 panel A). The two methods appear to produce estimates that are more similar to each other than to the ground truth labels (compare bottom row scatter plots in orange to the blue and red scatter plots in panel A). Fig. 2 panel B shows that the inferred median difference in APE, *µ*°*^d^*, between the method’s estimates for the three parameters is close to zero (*| µ*°*^d^ |* 95% HPI is *<* 4 ppts; SI Table S1; SI Figure S3).

We also compared the performance of uncertainty quantification using quantile-CNN-based conformalized quantile regression (CQR; Romano et al. 2019) to that of Bayesian HPIs for each of the experiments. We trained seven qCNNs to predict inner-quantiles at seven different levels to compare with the Bayesian HPIs; *τ* = *{*0.05, 0.1, 0.25, 0.5, 0.75, 0.9, 0.95*}*. We then used another simulated dataset to calibrate predicted intervals which we refer to as CPIs which theoretically have correct coverage properties (Romano et al. 2019) like the HPIs. For the test dataset of 138 trees, the CPIs had coverages that matched well with expectations to a comparable degree to the Bayesian HPI (Figure 2 panel C) though more variable. To further confirm that our CQR procedure was adequately calibrating the qCNN estimates, we confirmed correct coverages of CPIs for a much larger dataset with 5,000 trees (Figure 3). On average, the widths of CPIs in the set of 138 trees shown in (Figure 2) was about 20 - 40% wider than that of the corresponding HPI and Jaccard similarity index ranging from 0.66 to 0.75 suggesting a high degree of overlap between the intervals (SI Figure S4 and SI Table S2). These results indicate the probability level of the CPI, *e.g.* 95%, can be safely interpreted as the probability a parameter falls within the CPI. The wider intervals suggest the basic CQR method employed here is somewhat less precise and thus more conservative than the Bayesian method.

Our trained CNN provides nearly instantaneous estimates of model parameters. While the run time of the likelihood approach employed in this study scales linearly with the size of the tree, the neural network has virtually constant run times that are more than three orders of magnitude faster. Because simulation-trained neural networks have a one-time cost of simulating the training data set and then training the neural network, these methods are often called amortized-approximators (Bürkner et al. 2022). This means the time savings aren’t recouped until a certain number of trees have been analyzed. For example, here over 524 trees would need to be analyzed to realize the cost savings of simulating data and training our neural network (Figure 4). This illustrates the importance of simulation optimization and generality for likelihood-free approaches to inference.

**Figure 4:**
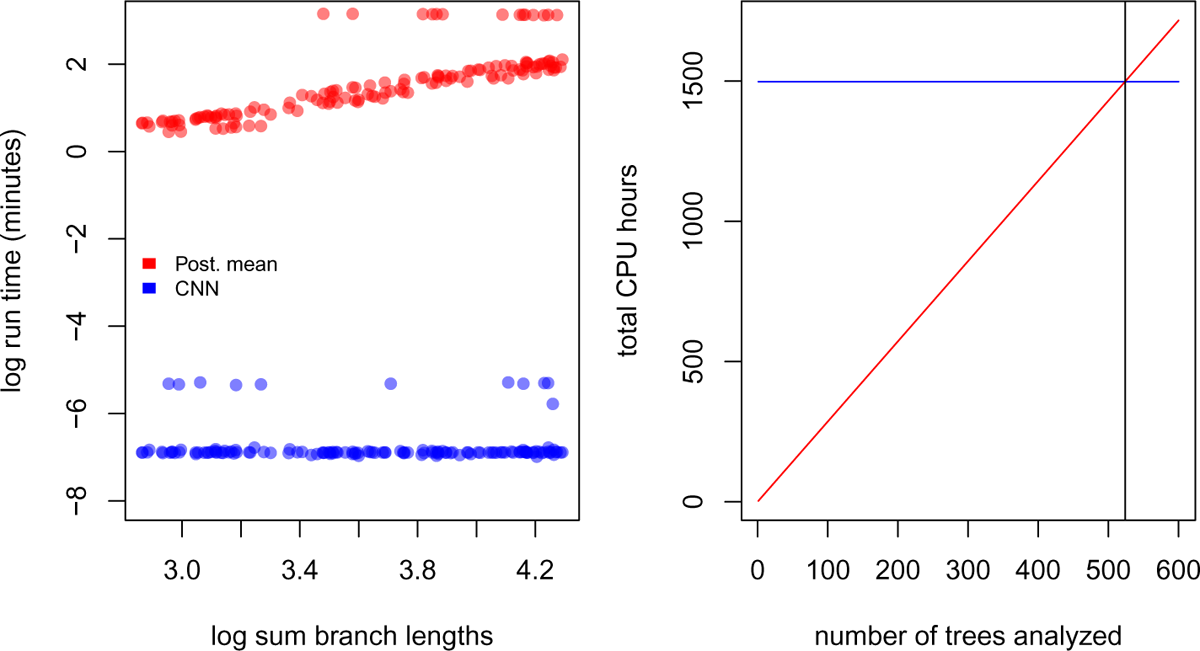
Left: Estimates of time to complete analysis of each of 138 trees relative to tree size. Right: The number of trees (524; gray vertical line) needed to analyze for total analysis time of Bayesian method (red line) to equal that of the entire simulation and CNN training and inference pipeline (blue line).

### Comparing sensitivity to model misspecification

To test the relative sensitivity of CNN estimates and the likelihood-based MPE to model misspecification, we simulated several test data sets under different, more complex epidemic scenarios and compared the decrease in accuracy (increase in APE).

Our first model misspecification experiment tested performance when assuming all locations had the same R_0_ when, in fact, each location had different R_0_*_i_* values. The median APE for all three parameters increased to varying degrees (SI Fig. S5 Panel A) compared to the median APE measured in Fig. S3. We found that both methods converged on similar biased estimates for R_0_. In both the CNN and Bayesian method, estimates of *δ* were relatively robust to misspecifying R_0_. In contrast, the migration rate showed much more sensitivity to this model violation in both methods with both methods also converging on similarly biased estimates (Figure 5 A). The median difference in error between the two methods is close to zero for all rate parameters (*| µ*°*^d^ |* 95% HPI *<* 6 ppts; SI Table S1) (SI Figure S5 Panel B). For both methods of uncertainty quantification the coverage declined by similar amounts for all three parameters with *δ* showing little to no sensitivity to R_0_ misspecification (Figure 5 panel C and SI Table S2). The patterns of coverage are also somewhat less regular across the qCNN quantiles than the HPIs for the migration rate parameter likely due in part to the fact that each inner quantile qCNN was trained independently and thus have independent errors. The relative interval widths and Jaccard similarity indexes did not change appreciably from predictions under the true model (SI Figure S4 and SI Table S2). Our CNN appears to be slightly more sensitive than the Bayesian approach when predicting the outbreak location. Nevertheless, their distributions are quite similar (Figure 5 Panel C).

**Figure 5:**
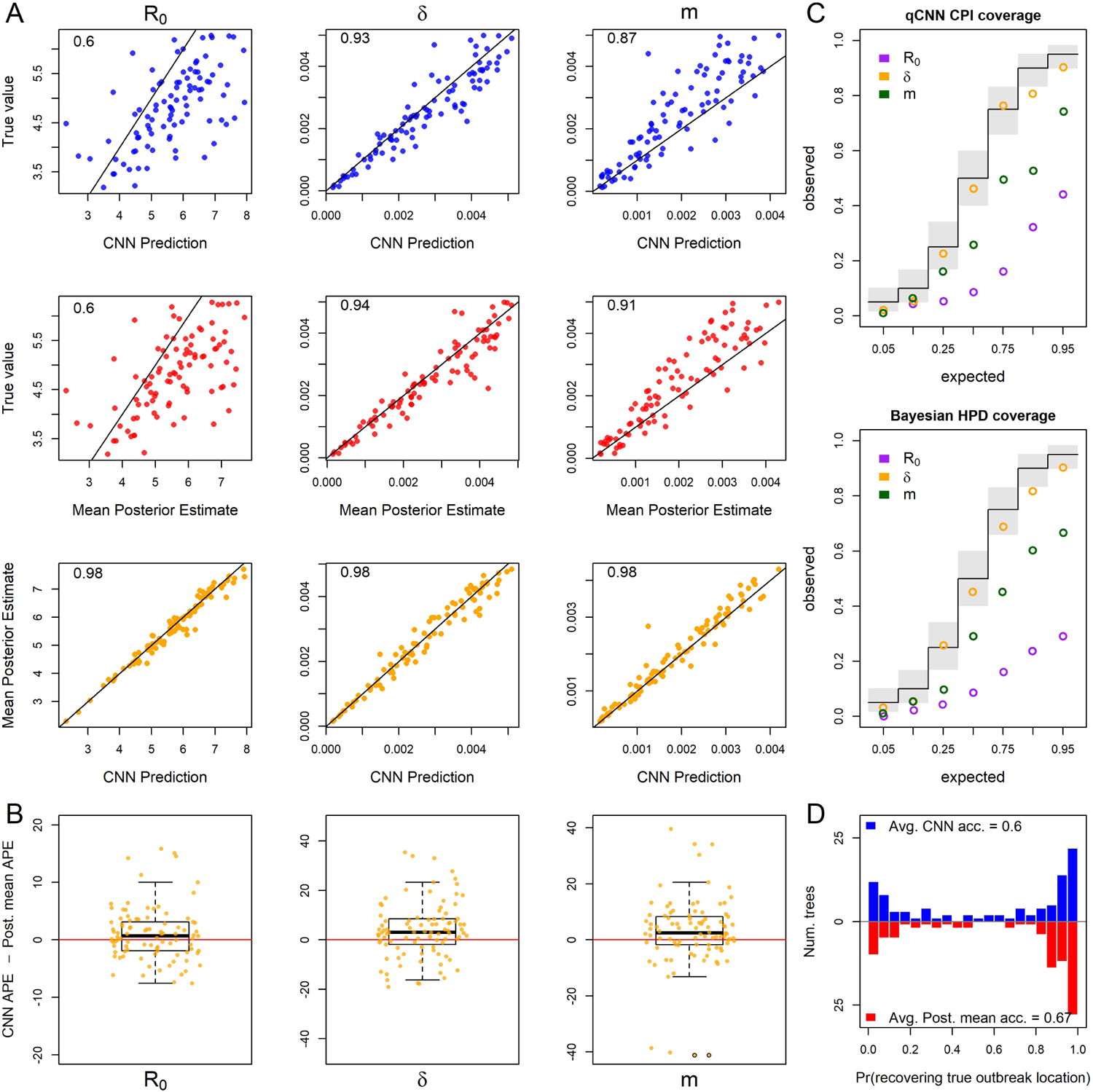
For 93 test trees where the R_0_ parameter was misspecified: the simulating model for the test data specified 5 unique R_0_s among the five locations while the inference methods assumed one R_0_ shared among locations. Because of this, the estimates for R_0_ are plotted against mean of the five true R_0_ values. See Figure 2 for general details about plots.

Next, we measured method sensitivity when the sampling process of the test trees violates assumptions in the inference model. In this set, each location had a unique and independent sampling rate, *δ*, rather than a single *δ* shared among locations. The median APE only increased for *δ* and m (SI Figure S7 Panel A). As expected, estimates of *δ* were highly biased for both methods (Figure 6 panel A). Panel A also shows that R_0_ is virtually insensitive to sampling model misspecification, but that migration rate, again, is highly sensitive in both the CNN and likelihood method. The median difference in error between the two methods is close to zero for all the rate parameters (*| µ*°*^d^ |* 95% HPI *<* 5 ppts; SI Table S1, SI Figure S7) (Figure 6 panel B). For both methods coverage declined for *δ* and m, while R_0_ showed little to no sensitivity to *δ* misspecification (Figure 6 panel C and SI Table S2). The relative widths and degree of overlap was again similar to the experiments above (SI Figure S8, SI Table S2). We again also see greater irregularity among CPI levels in coverage, notably *δ* at inner-quantile level 0.9. The location of outbreak prediction is also somewhat sensitive in both methods, with the CNN showing a slightly larger mean difference, but the overall distribution of accuracy of all the test trees again is similar (Figure 6 panel C).

**Figure 6:**
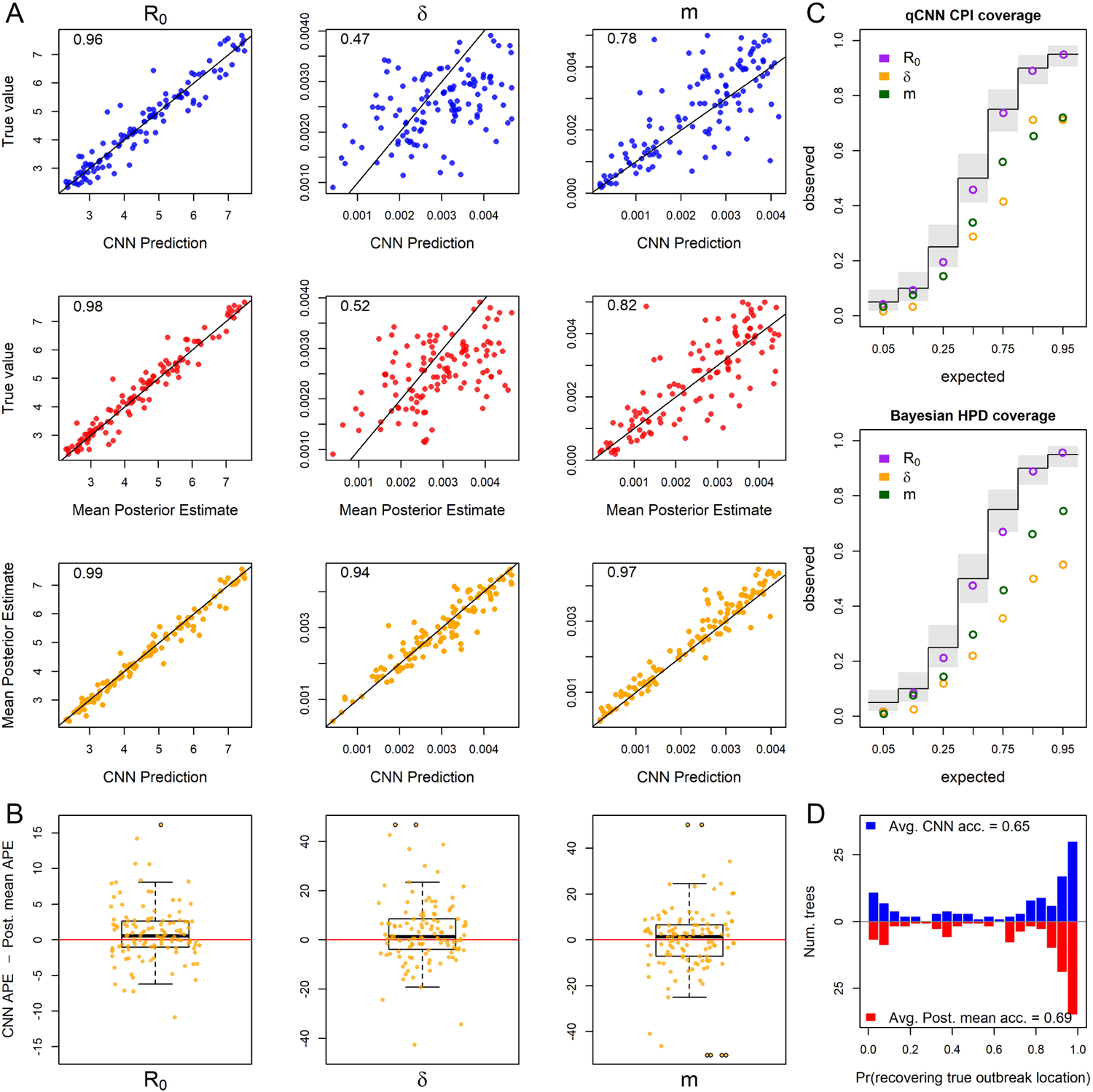
For 118 test trees where the sampling rate parameter was misspecified: the simulating model for the test data specified 5 unique sampling rates among the five locations while the inference methods assumed one sampling rate shared among locations. The estimates of *δ* are plotted against the mean true values of *δ*. See Figure 2 for general details about plots.

To explore sensitivity to migration model underspecification, we simulated a test set where the migration rates between locations is free to vary rather than being the same among locations as in the inference model. This implies 5! unique location-pairs and thus unique migration rates in the test data set. Results show that for both methods the parameters R_0_ and *δ* are highly robust to this simplification (SI Fig. S9 Panel A). Though estimates of a single migration rate had a high degree of error compared to a single pair of locations’ migration rates (Figure 7 panel A), the two methods still had similar estimates with the difference in APE centered near zero (Figure 7 panel B). The inferred median difference in APE was close to zero (*| µ*°*^d^ |* 95% HPI *<* 3 ppts; SI Table S1; SI Figure S9 Panel B). For both methods the coverage only declined significantly for the migration rate and the decrease was again similar in magnitude across quantiles (Figure 7 panel C and SI Table S2). Again, relative widths and degree of overlap of CPI and HPI were similar to previous experiments (SI Figure S10, SI Table S2) There was a slight but similar decrease in accuracy in predicting the outbreak location for both methods (Figure 7 panel C).

**Figure 7:**
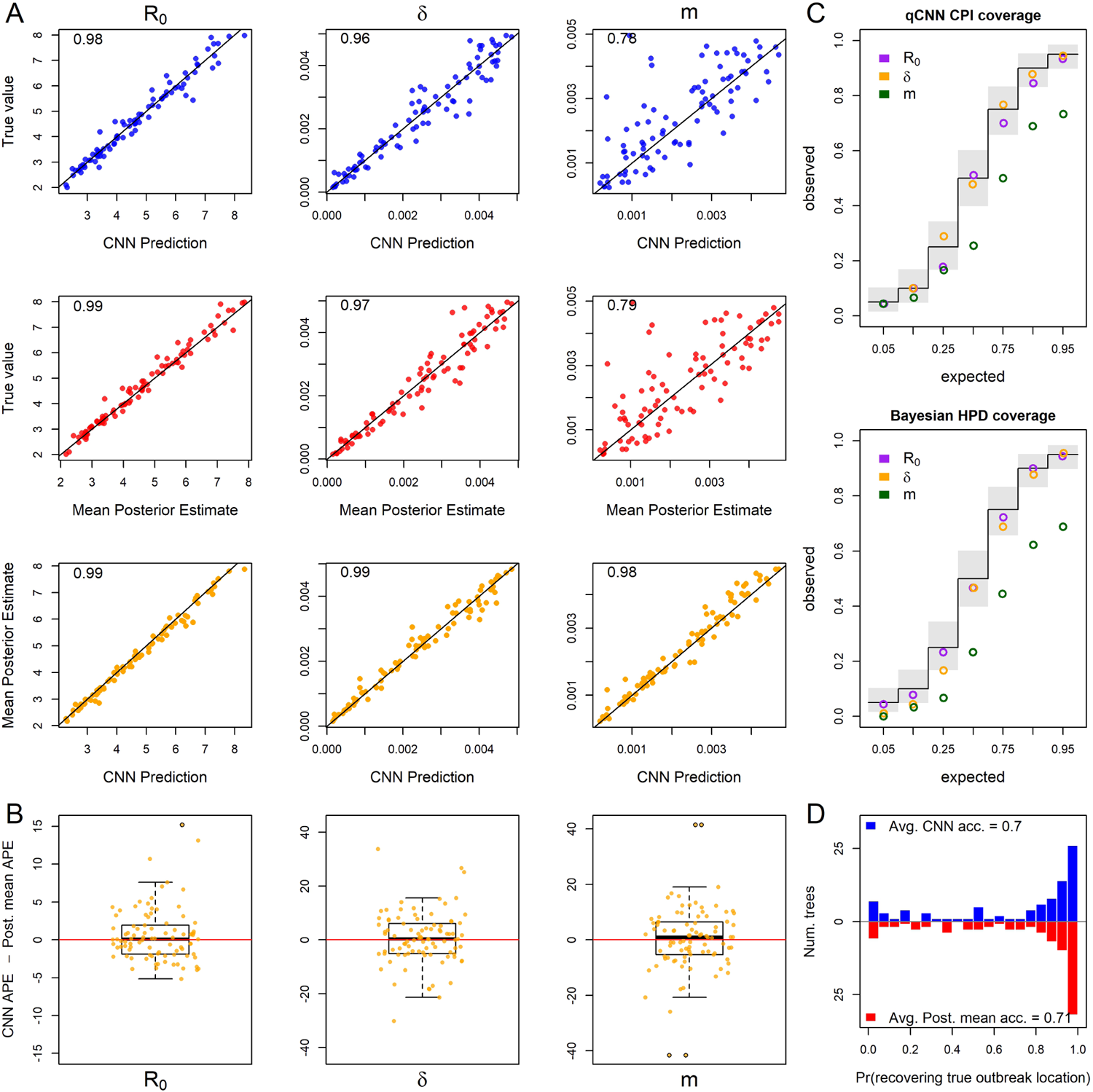
For 90 test trees where the migration rate parameter was misspecified: the simulating model for the test data specified 5! (120) unique migration rates among the unique pairs of the five locations while the inference methods assumed all migration rates were equal. The infered migration rate is plotted against the mean pairwise migraiton rates of test data set. See Figure 2 for general details about plots.

When testing the sensitivity of the two methods to arbitrary groupings of locations, we found that both methods showed equal sensitivity to the same parameters (Fig. 8 Panels A and B). In particular, the migration rate showed a modest increase in median APE and R_0_ and sample rate showed virtually no sensitivity to arbitrary grouping of locations (SI Figure S11 Panel A). The inferred median difference between method APE’s was again close to zero (*| µ*°*^d^ |* 95% HPI *<* 4 ppts; SI Table S1; SI Figure S11 Panel B). For both methods the coverage declined modestly only for the migration rate (Figure 5 panel C and SI Table S2). Relative widths and interval overlap showed virtually no change (SI Figure S12 and SI Table S1). These results suggest that for at least the exponential phase of outbreaks where rate parameters do not vary among locations, these models have a fair amount of robustness to the decisions leading to geographical division of continuous space into discrete space. The outbreak location showed higher accuracy in both methods due to the fact that the test data was no longer a flat distribution; the 6 combined locations should contain 60% of the outbreak locations (Figure 8 panel C).

**Figure 8:**
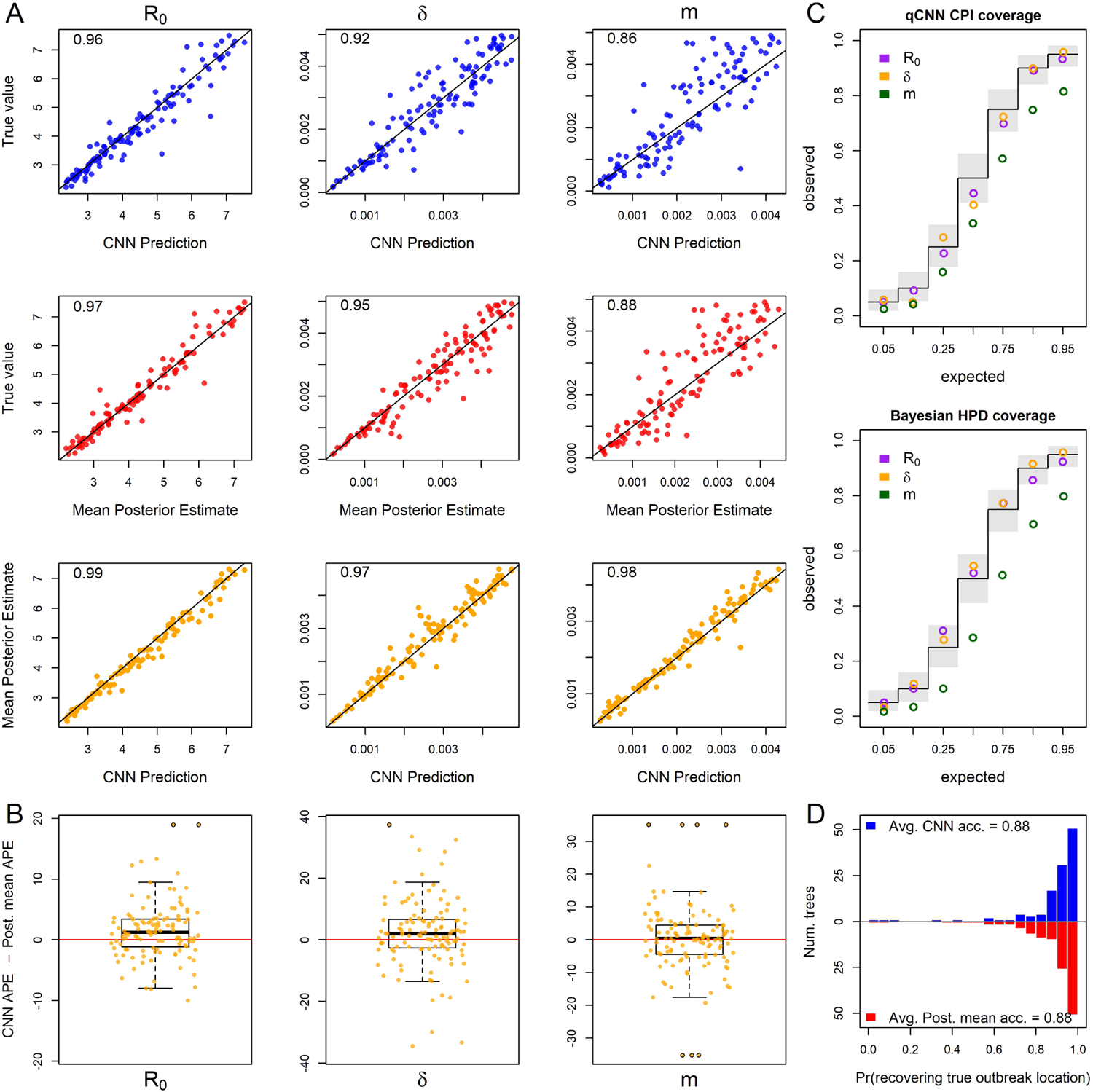
For 101 test trees where the number of locations was misspecified: the simulating model for the test data specified an outbreak among 10 locations with 6 locations subsequently combined into a single location while the inference methods assumed 5 locations with no arbitrary combining of locations. See Figure 2 for general details about plots.

Finally, we explored the relative sensitivity of our CNN to amounts of phylogenetic error that are present in typical phylogeographic analyses. Our simulated phylogenetic error produced trees with average Jaccard similarity indexes between the inferred tree and the true tree of about 0.5 with 95% of simulated trees having distances within 0.36 and 0.72.

We again compared inferences derived from the true tree and the tree with errors using the CNN and the Bayesian LIBDS methods. Results show that migration rate was minimally affected but R_0_ and *δ* were to a some degree sensitive to phylogenetic error (Figure 9 panel A; SI Figure S13 Panel A), with both methods again showing similar degrees of sensitivity (Figure 9 panel B). The inferred median difference was, yet again, small (*| µ*°*^d^ |* 95% HPI *<* 6 ppts. SI Table S1, SI Figure S13 Panel B). Coverages of *δ* declined for both methods in a similar way across quantiles. Again the 90% inner quantile showed some inconsistency with its nieghboring quantiles. In this case its coverage for *δ* was slightly higher than the 95th inner quantile. The CPIs for R_0_ appear much less sensitive (Figure 9 panel C and SI Table S2). Although the relative widths of the CPIs and HPIs were similar to previous experiments, the degree of overlap decreased somewhat by about 5 - 10% (SI Figure S14 and SI Table S2). One difference between this experiment and the others, is that trees are data instead of model parameters. It is interesting that the point estimates from the two methods show similar biases while the coverages seem to depart somewhat. Inference of the origin location, were very similar for both methods (Fig. 9 Panel C).

**Figure 9:**
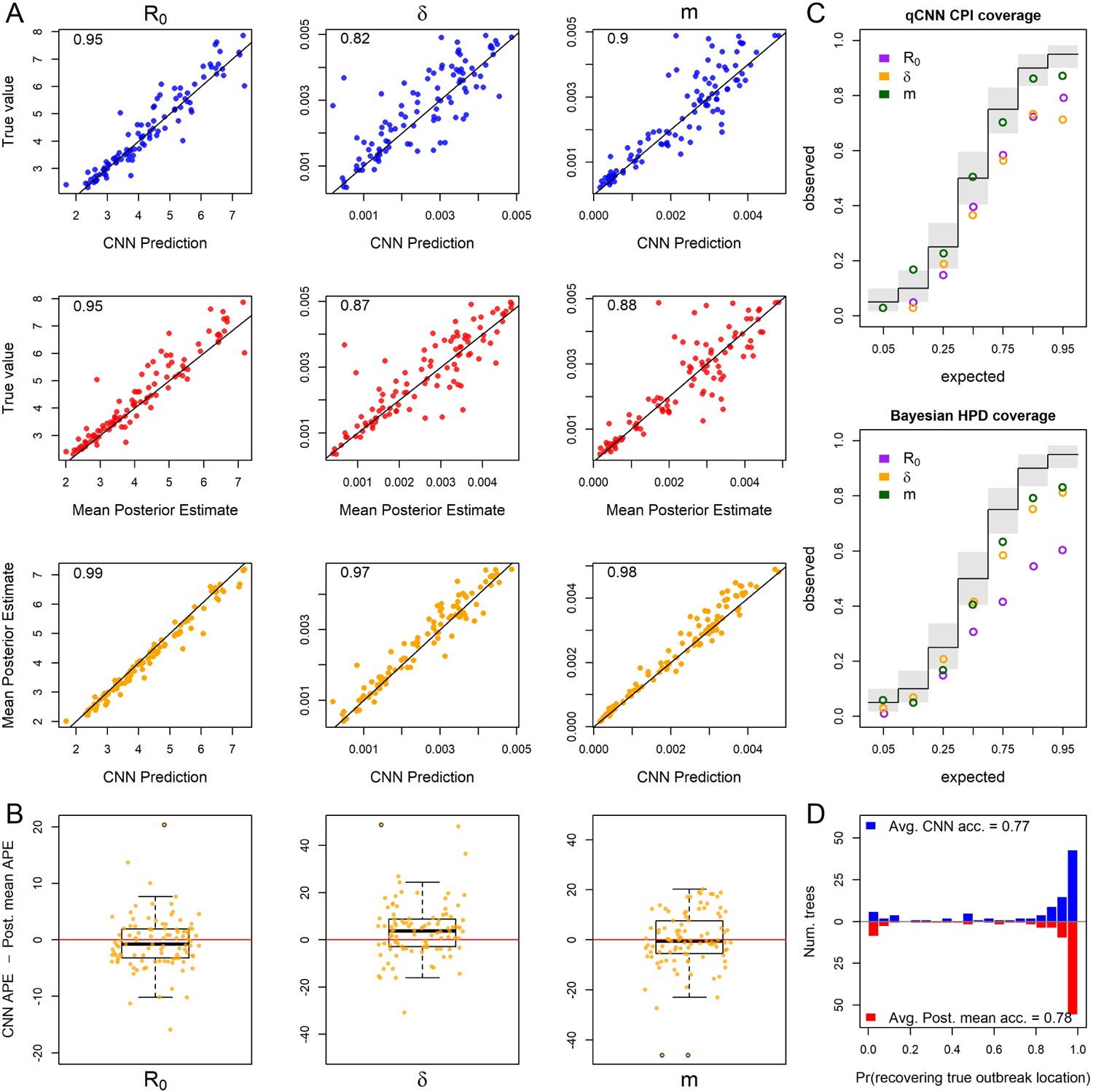
For 118 test trees where the time tree was misspecified: the true tree from the simulated test set was replaced with an inferred tree from simulated DNA alignments under the true tree. See Figure 2 for general details about plots.

### Analysis of SARS CoV-2 tree

We next compared our likelihood-free method to a recent study investigating the phylodynamics of the first wave of the SARS CoV-2 pandemic in Europe (Nadeau et al. 2021). Despite simulating the migration and the sampling processes differently from Nadeau et al. (2021), our CNN produces similar estimates for the location-specific R_0_ and the origin of the A2 clade (Figure 10). Whether the full tree or just the A2 clade is fed into the network, the predicted R_0_ for each location was not far from the posterior estimates of Nadeau et al. (2021). For the most part the R_0_ 95% CPI for each location overlaps to a high degree with the 95% HPI and is roughly 1.5 times wider indicating that our CNN estimates are relatively conservative. For Hubei the interval width of the a2 clade is much wider than the estimate using the whole tree. This is not surprising because there are no samples from Hubei in the a2 clade. We also obtained estimates for a single sampling rate and a single migration rate from our CNN and CPIs from our calibrated qCNN. Among the five location-specific estimates of the sampling proportion and the migration proportion from Nadeau et al. (2021), our CNN’s point estimates and interval estimates fall well within the their combined ranges.

**Figure 10:**
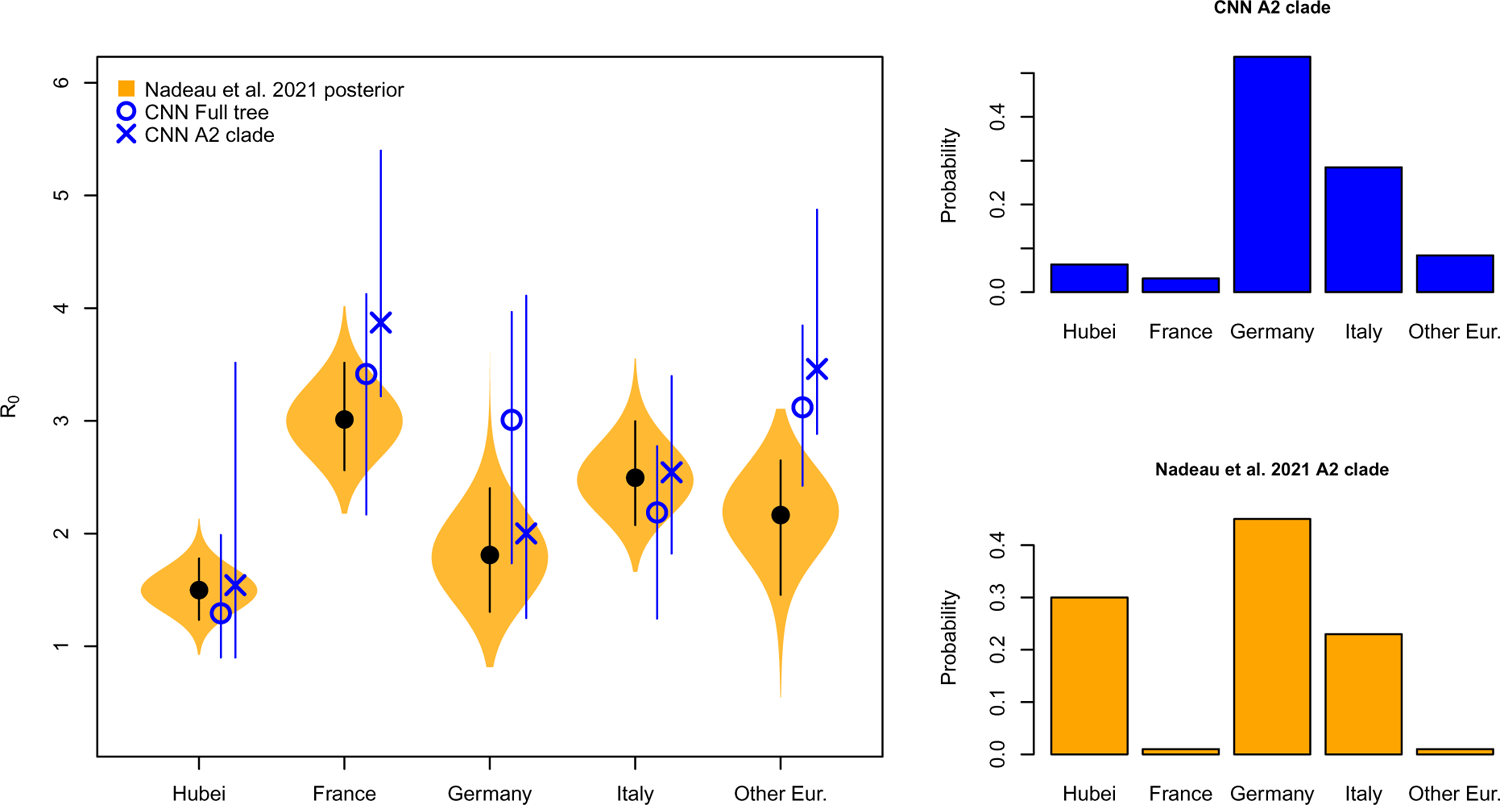
Location-dependent birth-death-sampling model (LDBDS) CNN comparison to (Nadeau et al. 2021) inference. Left violin plots show the posterior distributions of R_0_ for each location in Europe as well as Hubei, China (orange). The block dot and line within each violin plot shows the posterior mean and 95% HPI respectively. The blue X and O marks the LDBDS CNN prediction from analyzing the full tree and the A2 (European) clade respectively. Vertical blue lines give the 95% CPI for the CNN estimates of R_0_. Right barplots show the LDBDS CNN prediction (blue) and posterior inference (orange) from (Nadeau et al. 2021) of the ancestral location of the A2 (European) clade (see Figure 1 (Nadeau et al. 2021)).

The spillover location prediction CNN produced probability estimates of the A2 clade ancestral location the mostly agreed with that of Nadeau and colleagues (Figure 10, right histograms). The only significant discrepancy in the European origin prediction is that Nadeau and colleague’s analysis suggests a much higher probability that the most recent common ancestor of the A2 clade was in Hubei than our CNN predicts. This is likely because our CNN only used the A2 clade to predict A2 origins which has no Hubei samples to infer the origin of the A2 clade while Nadeau et al. (2021) used the whole tree. Notwithstanding this difference, among European locations, both methods predict Germany is the most likely location of the most recent common ancestor followed by Italy.

## Discussion and Conclusions

Inference models are necessarily simplified approximations of the real world. Both simulation-trained neural networks and likelihood-based inference approaches suffer from model under-specification and/or misspecification. When comparing inference methods it is important to assess the sensitivity of model inference to simplifying assumptions. In this study we show that newer deep learning approaches and standard Bayesian approaches behave and misbehave in similar ways under a panel of phylodynamic estimation tasks where the inference model is correct as well as when it is misspecified.

By extending new approaches to encode phylogenetic trees in a compact data structure (Voznica et al. 2022; Lambert et al. 2022), we have developed the first application of phylodynamic deep learning applied to phylogeography with serial sampling. Our approach is similar to that of Lambert et al. (2022) in which they analyzed a binary SSE model with exclusively extant sampling. By training a neural network on phylogenetic trees generated by simulated epidemics, we were able to accurately estimate key epidemiological parameters, such as the reproduction number and migration rate, in a fraction of the time it would take with likelihood-based methods. Like Voznica et al. (2022) and Lambert et al. (2022), we found that CNN estimators perform as well or nearly as well as likelihood-based estimators under conditions where the inference model is correctly specified to match the simulation model. The success of these separate applications of deep learning to different phylodynamic problems is a testament to the versatility of the CBLV encoding of trees.

We compared the sensitivity of deep learning and likelihood-based inference to model misspecification. Because deep-learning methods of phylogenetic and phylodynamic inference are new, few studies compare how simulation-trained deep learning methods fail in comparison to likelihood methods in this way (Flagel et al. 2019). We assume that when the inference model is correctly specified to match the simulation model, the trained CNN will, at best, produce noisy approximations of likelihood-based parameter estimates. In reality, issues related to training data set size, learning efficiency, and network overfitting may cause our CNN-based estimates to contain excess variance or bias when compared to Bayesian likelihood-based estimators. Our results from five model misspecification experiments show that both methods of inference perform similarly when the simulating model and the inference model assumptions do not perfectly match. These similarities exist not only in aggregate, when comparing method performance across datasets, but also when comparing performance for each individual dataset. This suggests that the CNN and likelihood methods are truly estimating parameters using isomorphic criteria, despite the fact that CNN heuristically learns these criteria through data patterns, while likelihood precisely and mathematically defines these criteria through the model definition itself.

Results of comparative sensitivity experiments like this are important because if likelihood-free methods using deep neural networks can easily be trained to yield estimates that are as robust to model misspecification as likelihood-based methods, then analysis of a large space of more complex outbreak scenarios for which tractable likelihood functions are not available can be developed and applied to real world data. Additionally, sufficiently realistic, pre-trained neural networks can yield nearly instantaneous inferences from data in real time to inform analysts and policy makers.

We also tested location-dependent SIR simulation trained neural network against a previous publication fitting a similar model – location-dependent birth-death-sampling (LDBDS) model – on real-world data using a Bayesian method. Our CNN predicted location-specific R_0_*_i_* and outbreak origin in Europe were similar to that inferred in (Nadeau et al. 2021). This result and our model misspecification experiments suggest that simulation-trained deep neural networks trained on phylogenetic trees can find patterns in the training data that generalize well beyond the training data set.

Our study extends the results of Voznica et al. (2022) and Lambert et al. (2022) in several important ways. Our work showed that the new compact bijective ladderized vector encoding of phylogenetic trees can easily be extended with one-hot encoding to include metadata about viral samples. Using this strategy, we trained a neural network to not only predict important epidemiological parameters such as R_0_*_i_* and the sampling rate, but also geographic parameters such as the migration rate and the location of outbreak origination or spillover. We anticipate that more diverse and complex metadata can be incorporated to train neural networks to make predictions about many important aspects of epidemiological spread such as the relative roles of different demographic groups and the overlap of different species’ ranges.

This approach can be readily applied to numerous compartment models used to describe the spread of different pathogens among different species, locations, and demographic groups, e.g. SEIR, SIRS, SIS, etc. (Ponciano and Capistŕan 2011; Volz and Siveroni 2018; Bjørnstad et al. 2020; Chang et al. 2020; O’Dea and Drake 2021) as well as modeling super-spreader dynamics as in (Voznica et al. 2022). Here we focused on one phase of outbreaks (the exponential phase), but there are many other scenarios to be investigated, such as when the stage of an epidemic differs among locations (e.g. exponential, peaked, declining). With likelihood-free methods, the link between the underlying population dynamics from which viral genomes are sampled and inferred phylogenetic trees can easily be interrogated. More complex models will require larger trees to infer model parameters. In this study we explored trees that contained fewer than 500 tips, but anticipate that larger trees will demonstrate even greater speed advantages of neural networks over likelhood-based methods either through subsampling regimes (Voznica et al. 2022) or by including larger trees in training datasets.

With fast, likelihood-free inference afforded by deep learning, the technical challenges shift from exploring models for which tractable likelihood functions can be derived towards models that produce realistic empirical data patterns, have parameters that control variation of those patterns, and are efficient enough to generate large training data sets. A growing number of advanced simulators are rapidly expanding the possibilities for deep learning in phylogenetics. For example, FAVITES (Moshiri et al. 2019) is a simulator of disease spread through large contact networks that tracks transmission trees and simulates sequence evolution. Gen3sis, MASTER, SLiM, and VGsim are flexible simulation engines for generating complex ecological, evolutionary, and disease transmission simulations (Hagen et al. 2021; Vaughan and Drummond 2013; Shchur et al. 2022; Haller and Messer 2019; Overcast et al. 2021). Continued advances in epidemic simulation speed and flexibility will be essential for likelihood-free methods to push the boundaries of epidemic modeling sophistication and usefulness.

There are several avenues of development still needed to realize the potential of likelihood-free inference in phylogeography using deep learning. The current setup is ideal for simulation experiments, but it is more difficult to ensure that the optimal parameter values for empirical data sets are within the range of training data parameters.

Standardizing input tree height, geographical distance, and other parameters help make training data more universally applicable. Simulation-trained neural networks are often called amortized methods (Bürkner et al. 2022; Schmitt et al. 2022) because the cost of inference is front-loaded, *i.e.* it takes time to simulate a training set and train a neural network. The total cost in time per phylogenetic tree amortizes as the number of trees analyzed by the trained model increases. These methods are therefore important when a model is intended to be widely deployed or be responsive to an emerging outbreak where policy decisions must be formulated rapidly. Because amortized approximate methods require multiple analyses to realize time savings, researchers need to generate training data sets over a broad parameter and model space so that trained networks can be applied to new and diverse data sets.

Our analysis introduces a simple approach to estimate the ancestral state corresponding to the root node or stem node of a phylogeny. More sophisticated supervised learning approaches will be needed to train neural networks to predict the ancestral locations for internal nodes other than the root. The topologies and branch lengths of random phylogenies in the training and test datasets will vary from tree to tree. Our approach relies on the fact that all trees contain a root node, meaning all trees can help predict the root node’s state. However, few (if any) trees in the training dataset will contain an arbitrary clade of interest within a test dataset, suggesting to us that naive approaches to train networks to estimate ancestral states for all internal nodes will probably fail. We are unaware of any existing solutions for generalized ancestral state estimation using deep learning, and expect the problem will gather more attention as the field matures.

Quantifying uncertainty is crucial to data analysis and decision making, and Bayesian statistics provides a framework for doing so in a rigorous way. It is essential to understand how uncertainty estimation with likelihood-free methods compare to likelihood-based methods when confronted with the mismatch of models and real-world data-generating processes. We quantified uncertainty using conformalized quantile regression (CQR; Romano et al. 2019) by training neural networks to predict quantiles and then calibrating those quantiles to produce the expected coverage. We refer to the resulting intervals as CPI and demonstrate that they predict well the coverage of true values on a test dataset (Figure 3) and behave in similar ways to Bayesian methods when the model is or is not misspecified (Figures 2 - 9). Despite having the same (correct) coverage as the Bayesian HPI, the interval length was 20-50% wider on average making them a more conservative (less precise) estimation procedure. Though this can likely be improved with more training data for qCNNs, there are more fundamental challenges for uncertainty quantification with quantile regression and conformalization.

Methods for estimating more precise intervals is an active vein of research among machine learning researchers and statisticians (Barber et al. 2020; Chung et al. 2021; Sousa et al. 2022; Gibbs et al. 2023). For example, although intervals estimated by the qCNN are conditional on each data point, the calibration of quantiles through CQR involves estimating marginal calibration terms that shift all quantiles by the same amount. If the error in the quantile coverage is not constant across the prediction range, then a more adaptive procedure should yeild more precise intervals (Sousa et al. 2022; Gibbs et al. 2023).

We also compared the consistency among CPI estimates at different inner-quantiles to that of HPIs at those same quantiles. We find that independently trained neural networks for each *α* level can potentially lead to inconsistencies where narrower, nested inner quantiles can have close to or higher coverage than wider quantiles (*e.g.* Figure 9 C).

Overall, our results suggest CQR is approximately consistent with likelihood-based methods and similarly sensitive to model misspecification, while there is room for improvement. Methods where all quantiles of interest can be estimated jointly (Chung et al. 2021) may be a fruitful avenue of research for such improvements.

Another important challenge of inference with deep learning is the problem of convergence to a location on the loss function surface that approximates the maximum likelihood well. There are a number of basic heuristics that can help such as learning curves but more rigorous methods of ascertaining convergence is the subject of active research (Bürkner et al. 2022; Schmitt et al. 2022).

With recent advances in deep learning in epidemiology, evolution, and ecology (Battey et al. 2020; Schrider and Kern 2018; Voznica et al. 2022; Radev et al. 2021; Lambert et al. 2022; Rosenzweig et al. 2022; Suvorov and Schrider 2022) biologists can now explore the behavior of entire classes of stochastic branching models that are biologically interesting but mathematically or statistically prohibitive for use with traditional likelihood-based inference techniques. Beyond epidemiology, we anticipate that deep learning approaches will be useful for a wide range of currently intractable phylogenetic modeling problems. Many phylogenetic scenarios – such as the adaptive radiation of anoles (**?**) or the global spread of the grasses (**?**) – involve the evolution of discrete traits, continuous traits, speciation, and extinction within an ecological or spatial context across a set of co-evolving species. Deriving fully mechanistic yet tractable phylogenetic model likelihoods for such complex scenarios is difficult, if not impossible. Careful development and applications of likelihood-free modeling methods might bring these phylogenetic scenarios into renewed focus for more detailed study. Although we are cautiously optimistic about the future of deep learning methods for phylogenetics, it will become increasingly important for the field to diagnose the conditions where phylogenetic deep learning underperforms relative to likelihood-based approaches, and to devise general solutions to benefit the field.

## Acknowledgements

We are grateful to Jacob McCord, Mark Lowell, Michael May, Fábio Mendes, Sarah Swiston, Sean McHugh, Walker Sexton, and Mariana Braga for helpful comments on the research.

## Data available from the Dryad Digital Repository

https://doi.org/10.25338/B8SH2J (Thompson et al. 2023) and code is available on github: https://github.com/ammonthompson/phylogeoepicnn

## Supplemental Tables

**Table S1:**
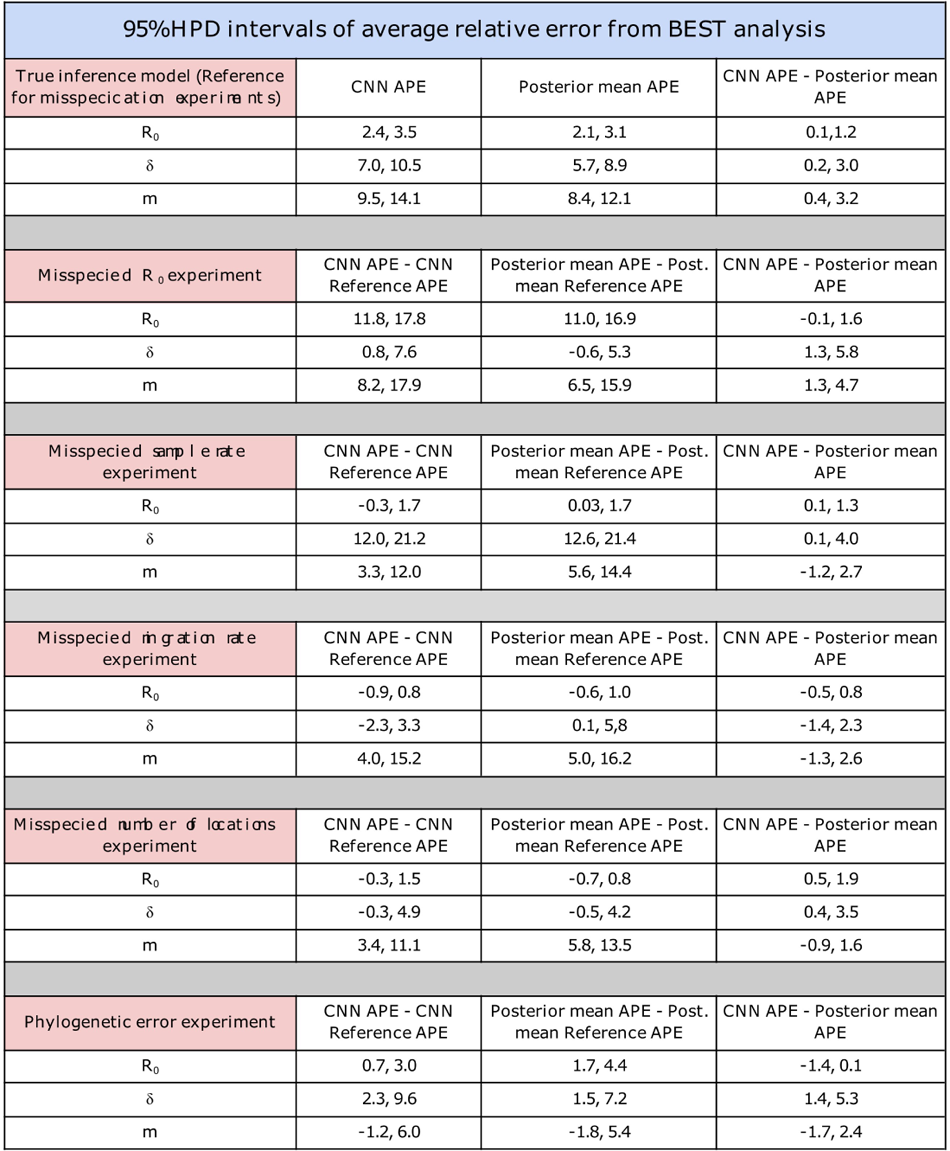
BEST comparisons between CNN and Bayesian absolute percent errors (APEs) for model parameters across all experiments.

**Table S2:** Comparison 95% CPI and HPI for all experiments.

| Coverage,<br>width, and<br>overlap of<br>95%<br>Intervals | $R_0$ | | | | $\delta$ | | | | $m$ | | | |
| --- | --- | --- | --- | --- | --- | --- | --- | --- | --- | --- | --- | --- |
|  | CNN<br>CPI | Bayes<br>HPI | Mean<br>CPI<br>width /<br>HPI<br>width | Mean<br>Jaccard<br>index | CNN<br>CPI | Bayes<br>HPI | Mean<br>CPI<br>width /<br>HPI<br>width | Mean<br>Jaccard<br>index | CNN<br>CPI | Bayes<br>HPI | Mean<br>CPI<br>width /<br>HPI<br>width | Mean<br>Jaccard<br>index |
| True model | 0.95 | 0.96 | 1.4 | 0.67 | 0.96 | 0.94 | 1.4 | 0.66 | 0.94 | 0.95 | 1.2 | 0.75 |
| Misspecified $R_0$ | 0.44 | 0.29 | 1.5 | 0.63 | 0.9 | 0.90 | 1.5 | 0.63 | 0.74 | 0.67 | 1.2 | 0.76 |
| Misspecified $\delta$ | 0.95 | 0.96 | 1.4 | 0.67 | 0.71 | 0.55 | 1.3 | 0.69 | 0.72 | 0.75 | 1.2 | 0.75 |
| Misspecified $m$ | 0.93 | 0.94 | 1.5 | 0.63 | 0.94 | 0.96 | 1.5 | 0.65 | 0.73 | 0.69 | 1.3 | 0.73 |
| Misspecified.<br>Number of<br>locations | 0.93 | 0.92 | 1.4 | 0.65 | 0.96 | 0.96 | 1.4 | 0.68 | 0.82 | 0.80 | 1.2 | 0.76 |
| Phylogenetic<br>error | 0.79 | 0.60 | 1.4 | 0.59 | 0.71 | 0.81 | 1.3 | 0.59 | 0.87 | 0.83 | 1.3 | 0.71 |

## Supplemental Figures

**Figure S1:**
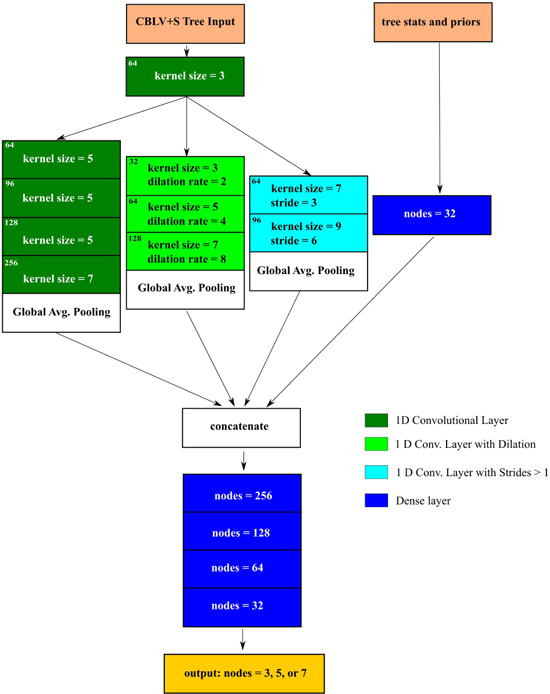
Diagram of deep neural network trained to make 2 kinds of predictions (rates and origin location) under two models (LIBDS and LDBDS).

**Figure S2:**
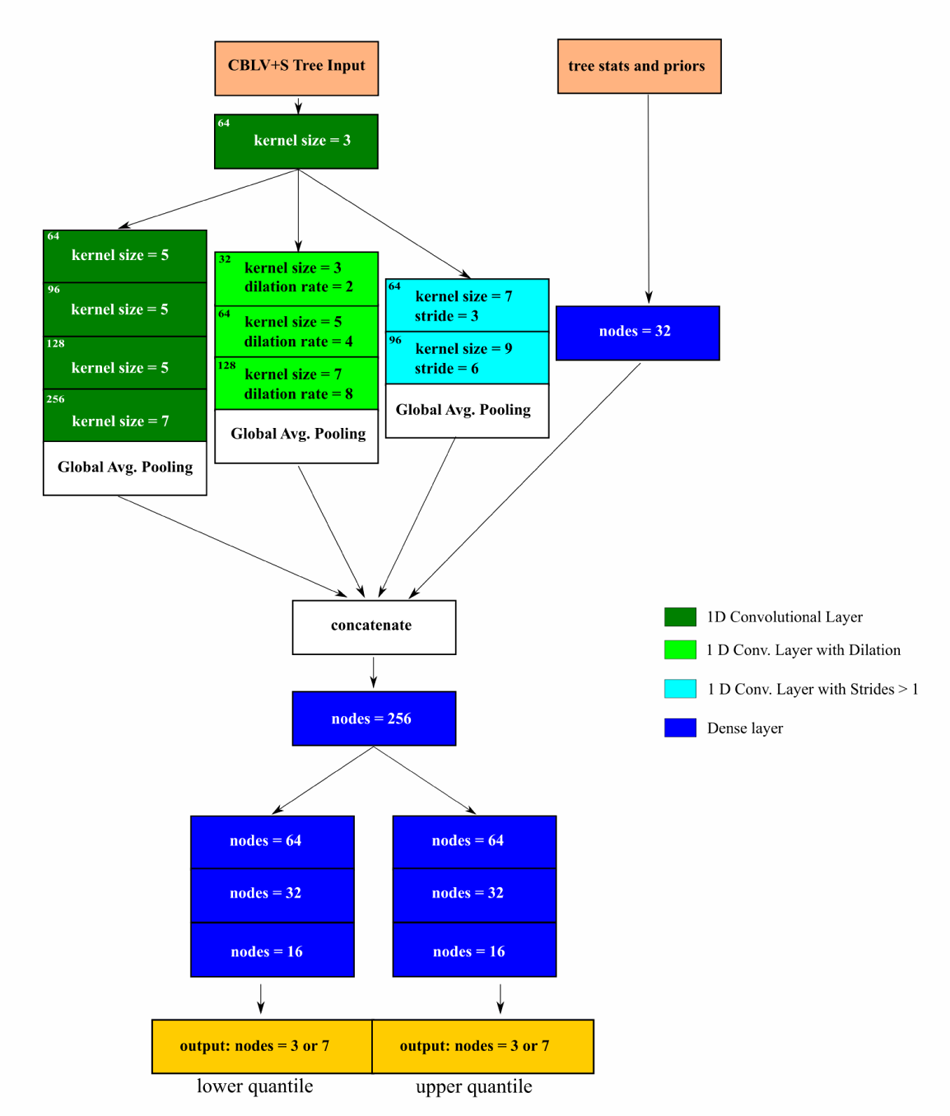
Diagram of deep neural network trained to predict the upper and lower quantiles for a specified *α* level under two models (LIBDS and LDBDS).

**Figure S3:**
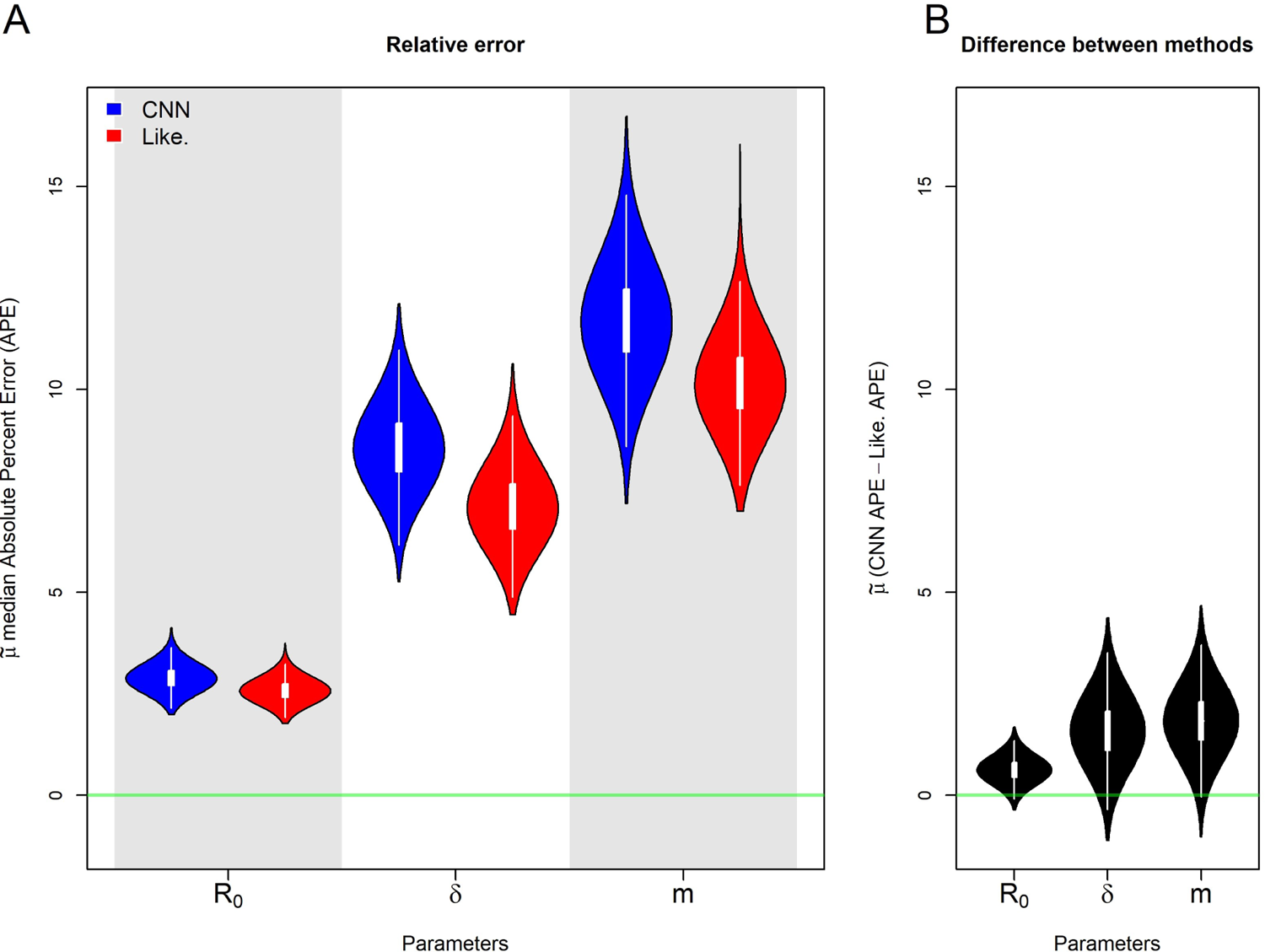
Posterior distributions of the population median, *µ*°, APE estimates of the rate parameters R_0_, *δ*, and m under the true model. A) shows posterior distribution of the median APE for each of the 3 rate parameters estimated by the CNN (blue) and the likelihood-based method (red). The green line indicates no error. B) shows the posterior distribution for the median difference between the CNN estimate’s APE and the likelihood-based estimate’s APE. The green line indicates the median APE difference is zero.

**Figure S4:**
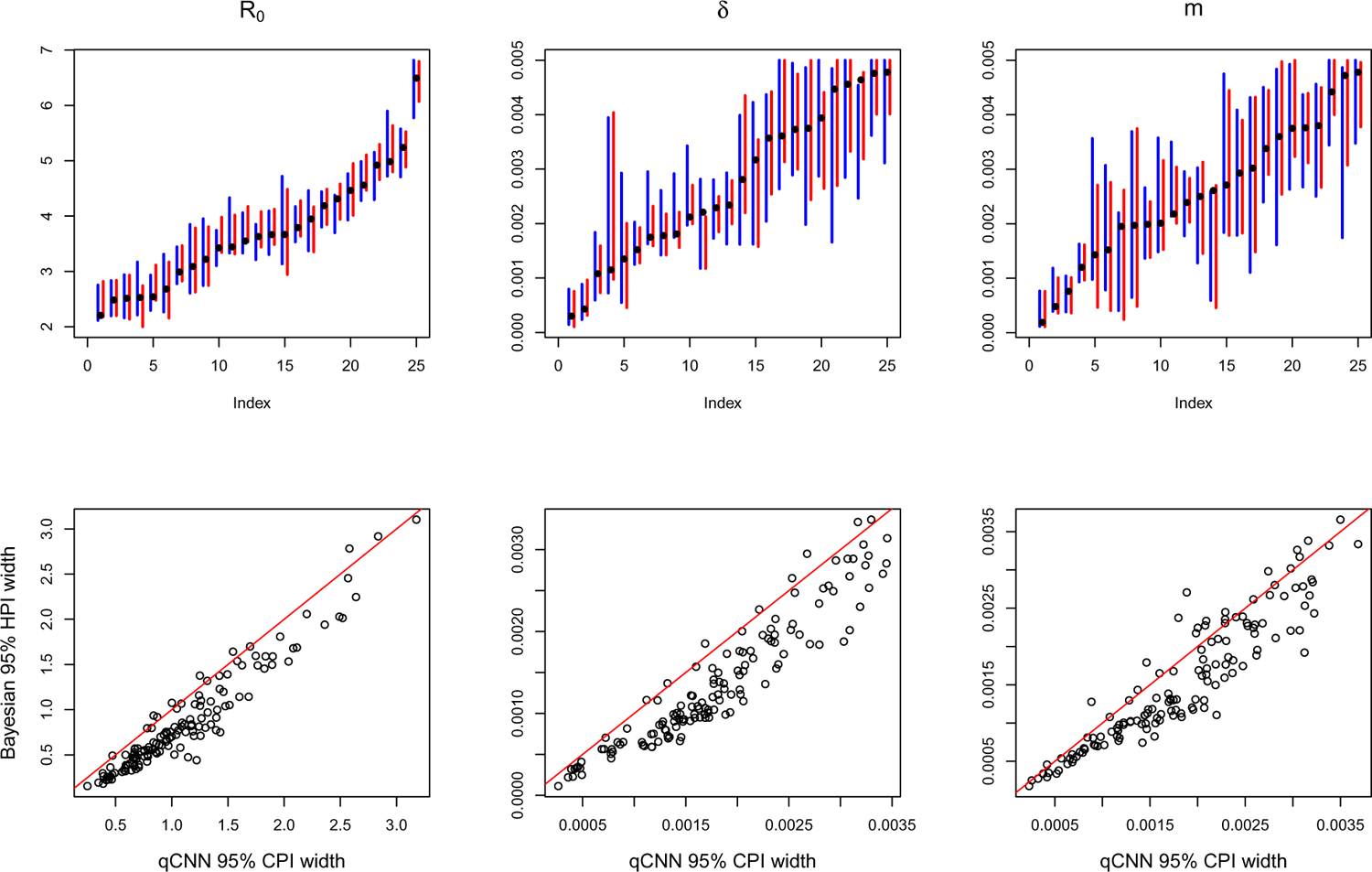
Comparison of interval overlap and relative widths of qCNN and Bayesian methods of uncertainty quantification under the true simulating model. Top row: 95% CPI from CNN conformalized quantile regression (blue). and 95% HPI from Bayesian phylogenetic analysis (red) from a random subset of the data for visualization purposes. Bottom row: scatterplots of the lengths of CPI and HPI intervals of all experiment data. The red diagonal y = x line is for reference.

**Figure S5:**
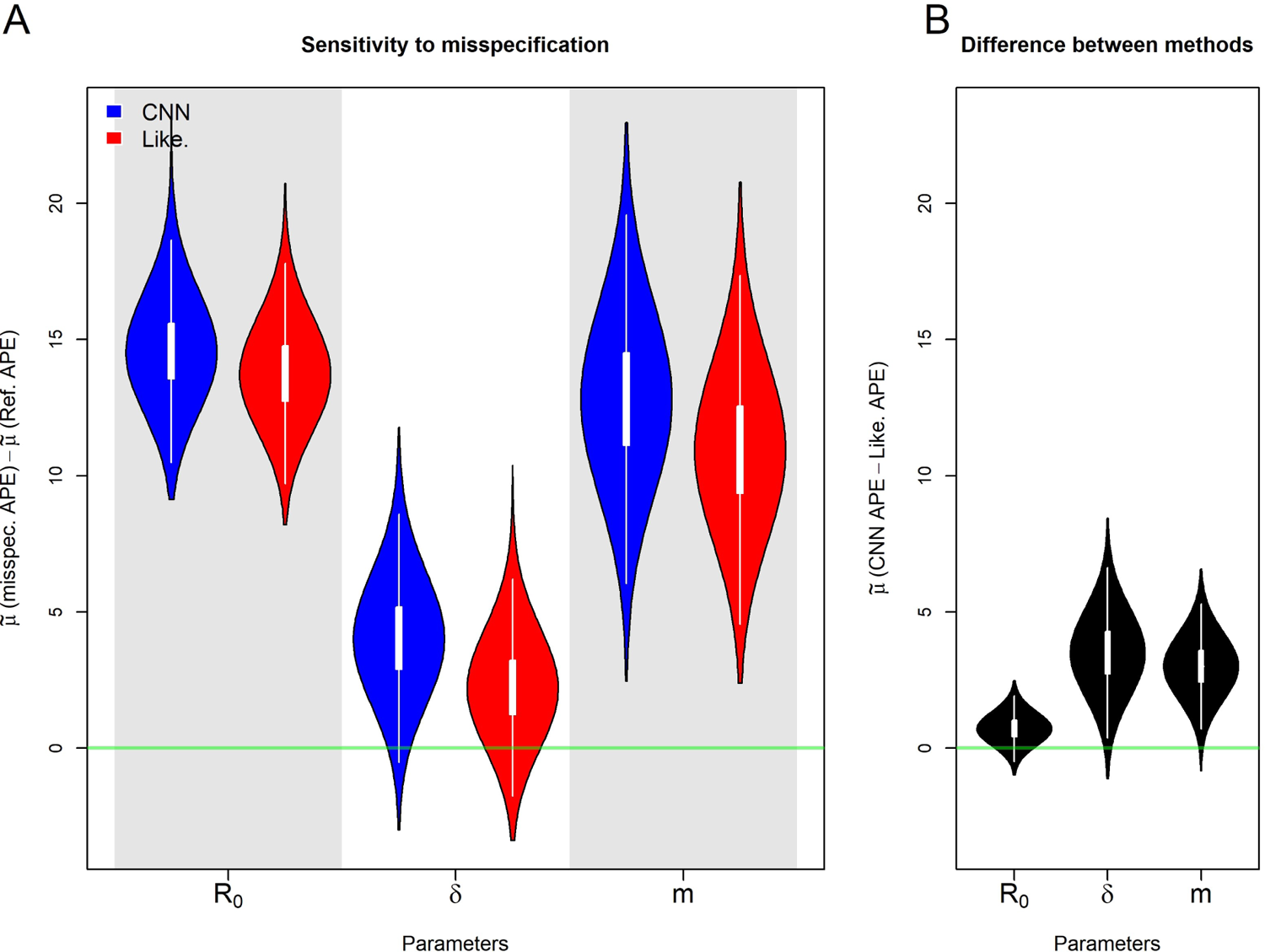
Posterior distributions of median, *µ*°, APE for the misspecified R_0_ experiment. A) shows posterior distribution of the difference between the median error under the misspecified model and the the median error under the true, reference model. B) shows the posterior distribution for the population median difference between the CNN estimate’s APE and the likelihood-based estimate’s APE.

**Figure S6:**
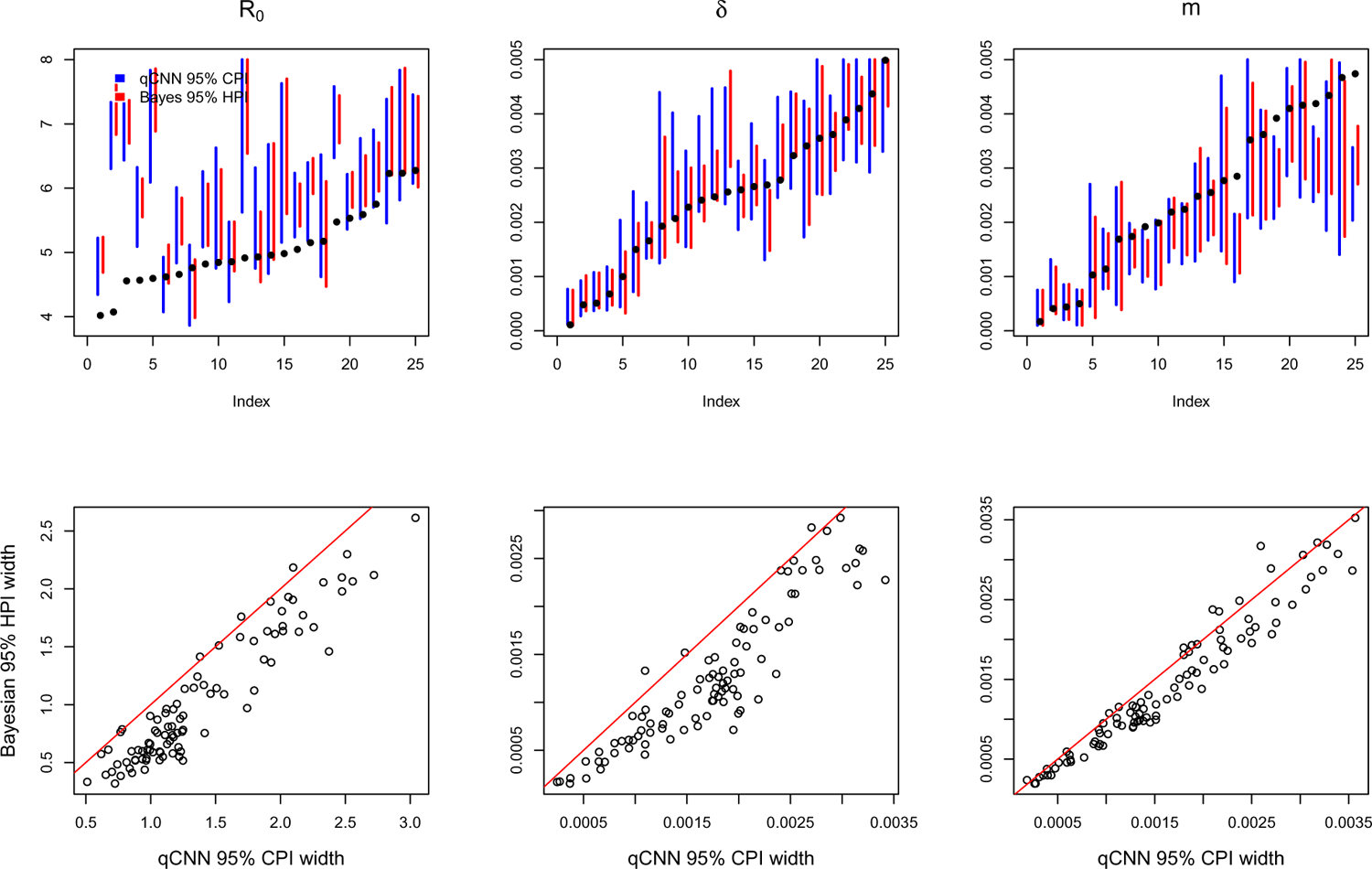
Comparison of CPI and HPI intervals for misspecified R_0_ experiment. See SI Figure S4 for general details about plot.

**Figure S7:**
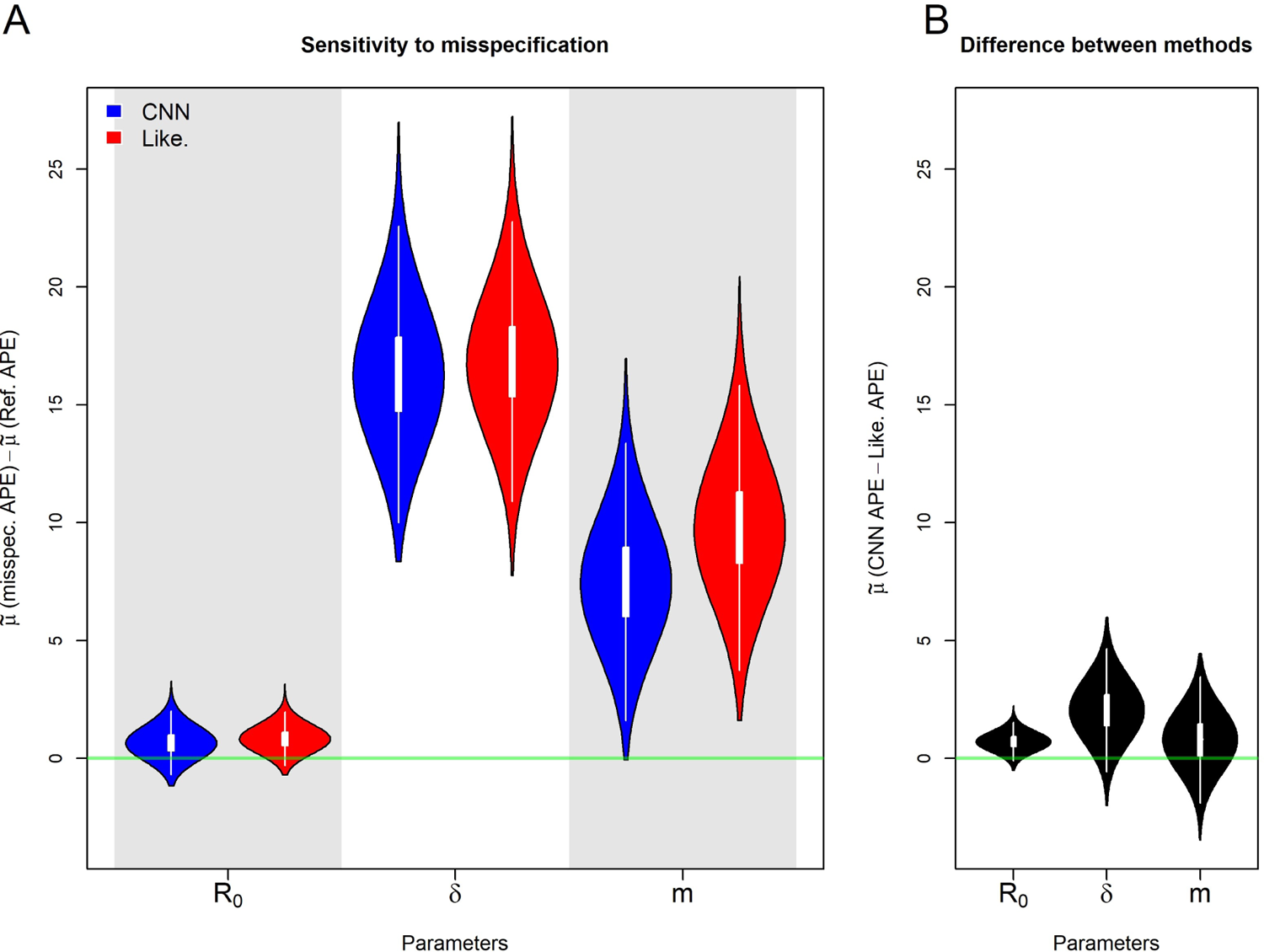
Posterior distributions of median, *µ*°, APE for the misspecified sampling rate, *δ*, experiment. Details are the same as in S5

**Figure S8:**
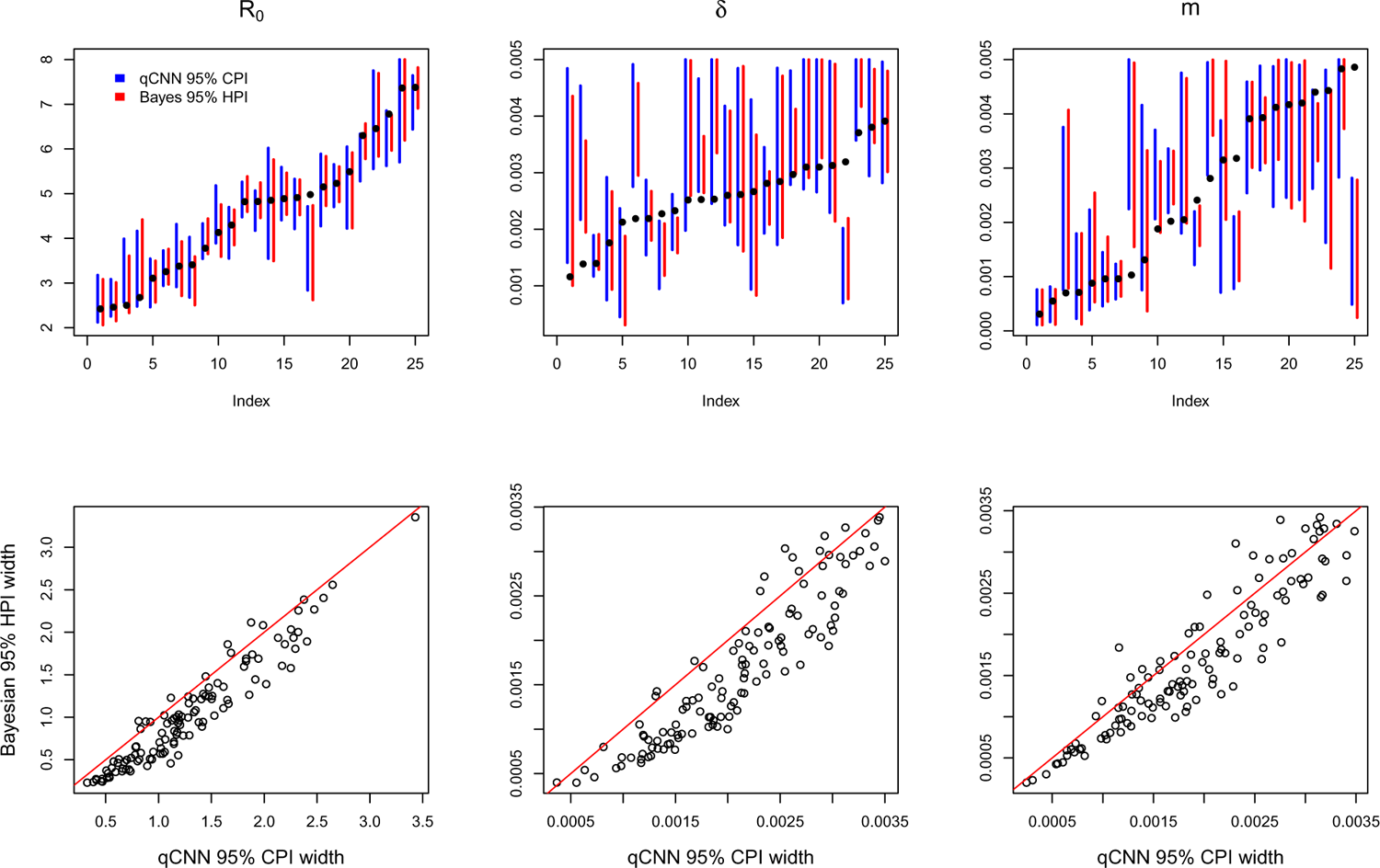
Comparison of CPI and HPI intervals for misspecified *δ* experiment. See SI Figure S4 for general details about plot.

**Figure S9:**
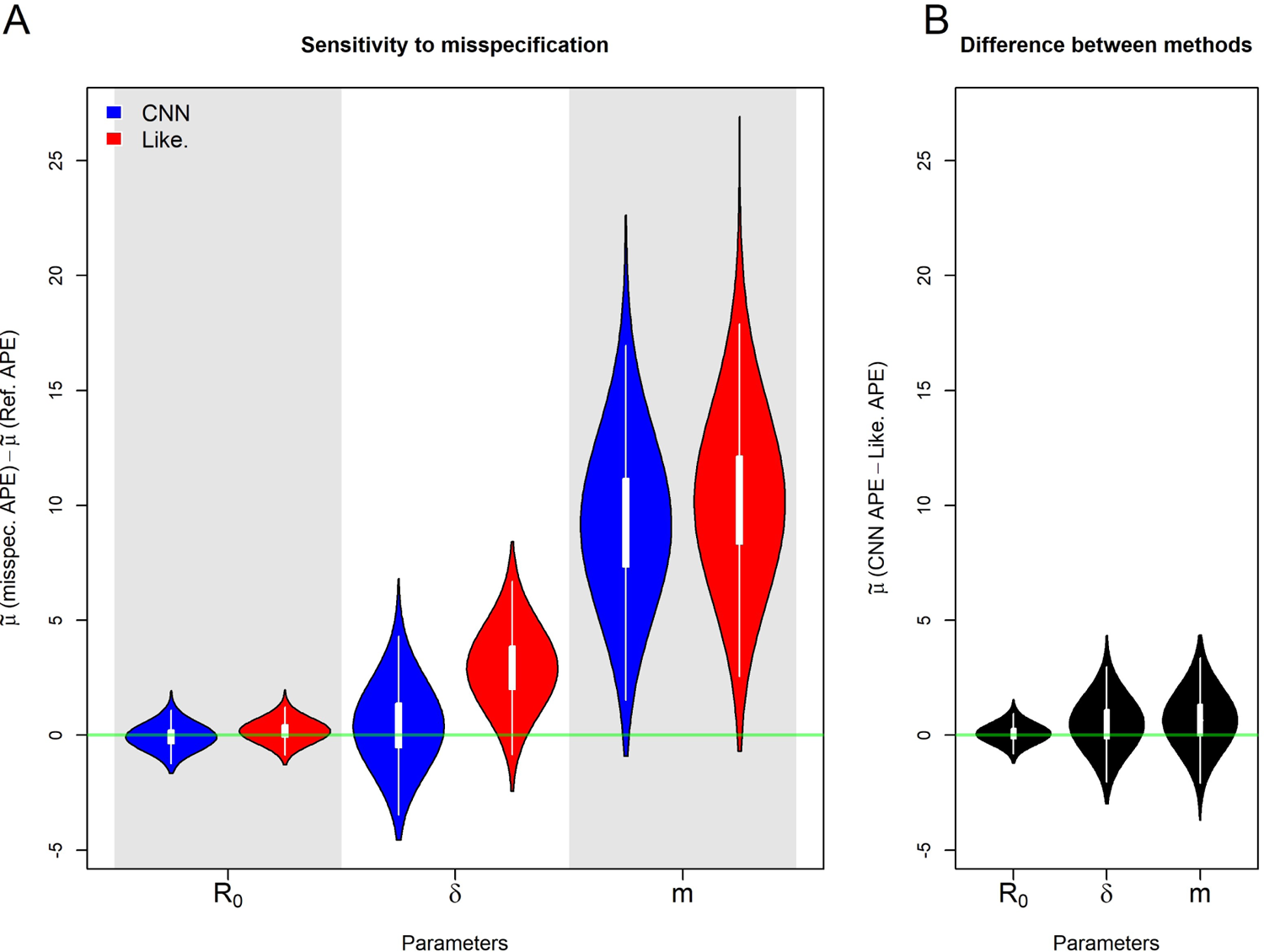
Posterior distributions of median, *µ*°, APE for the misspecified migration rate, m, experiment. Details are the same as in S5

**Figure S10:**
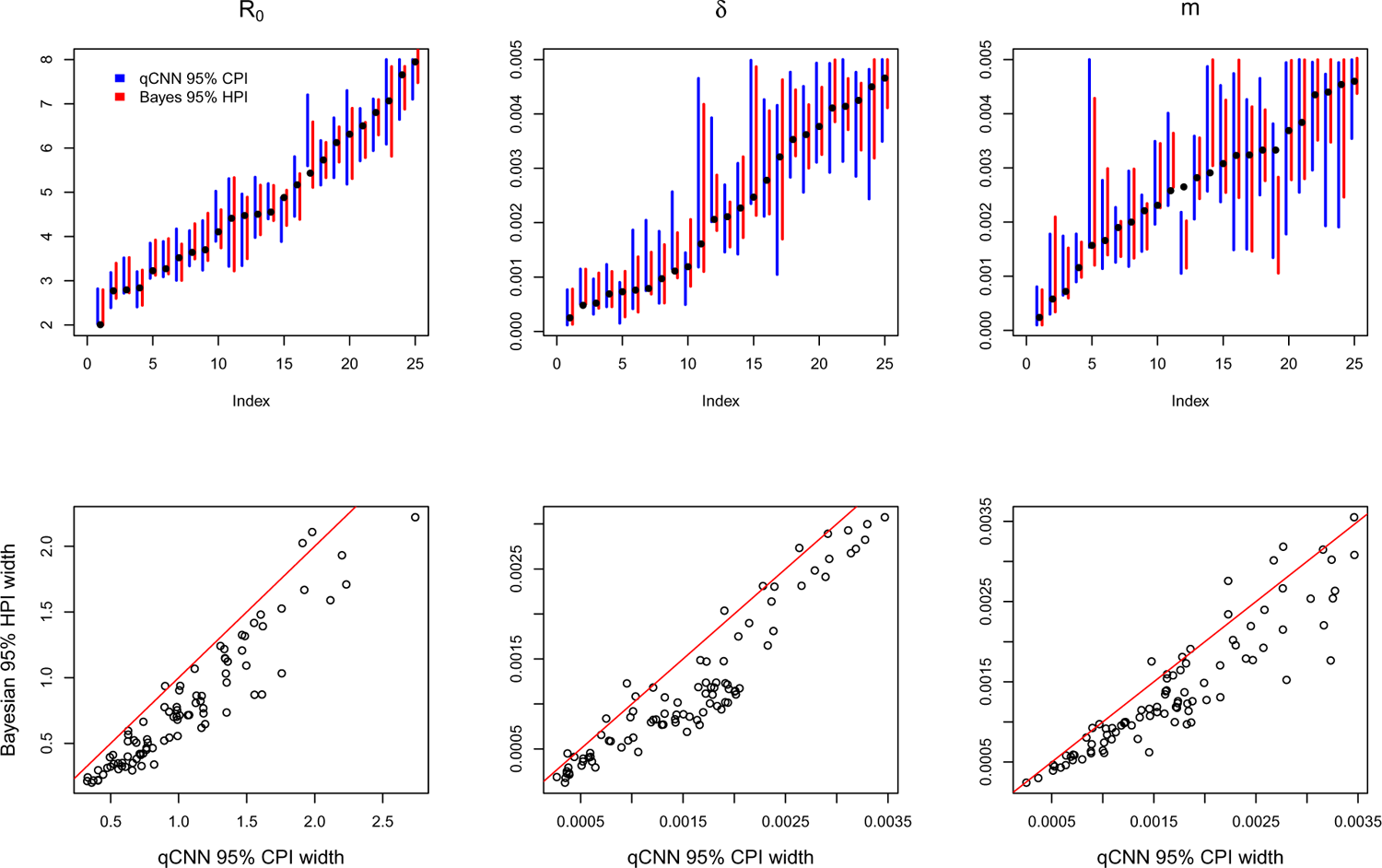
Comparison of CPI and HPI intervals for misspecified migration rate experiment. See SI Figure S4 for general details about plot.

**Figure S11:**
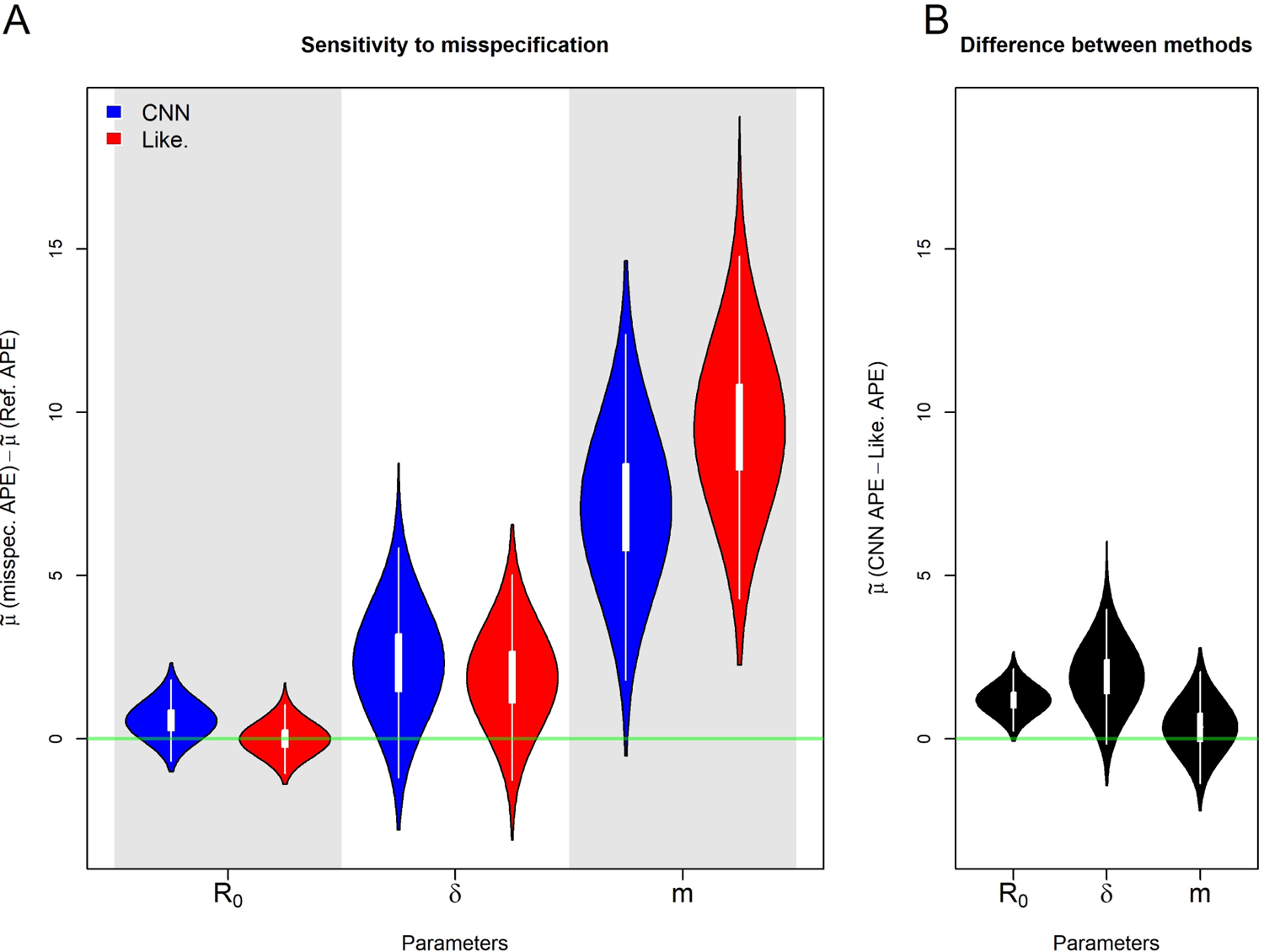
Posterior distributions of the median APE when the model is misspecified for the number of locations. Details are the same as in S5

**Figure S12:**
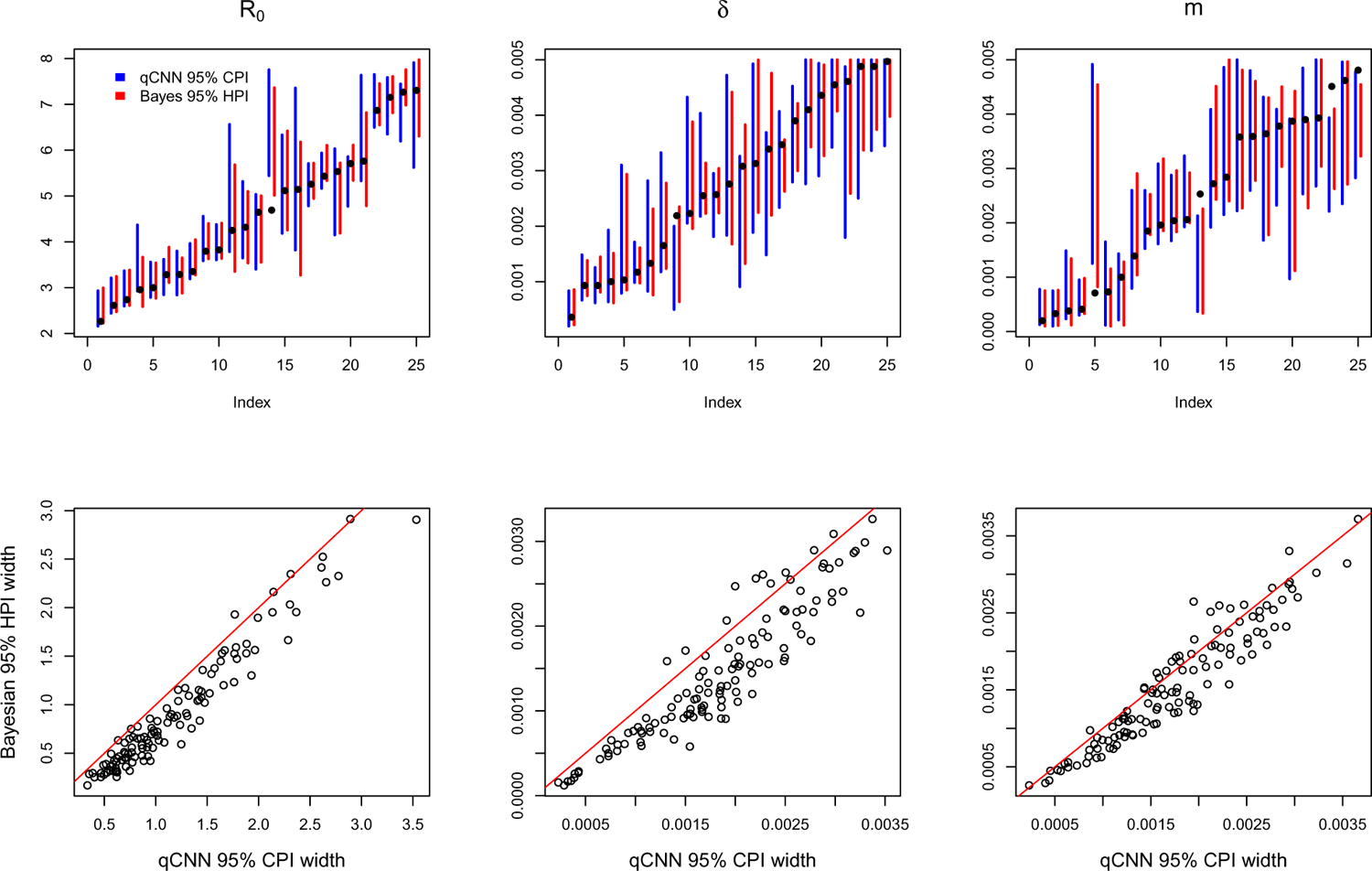
Comparison of CPI and HPI intervals for misspecified number of locations experiment. See SI Figure S4 for general details about plot.

**Figure S13:**
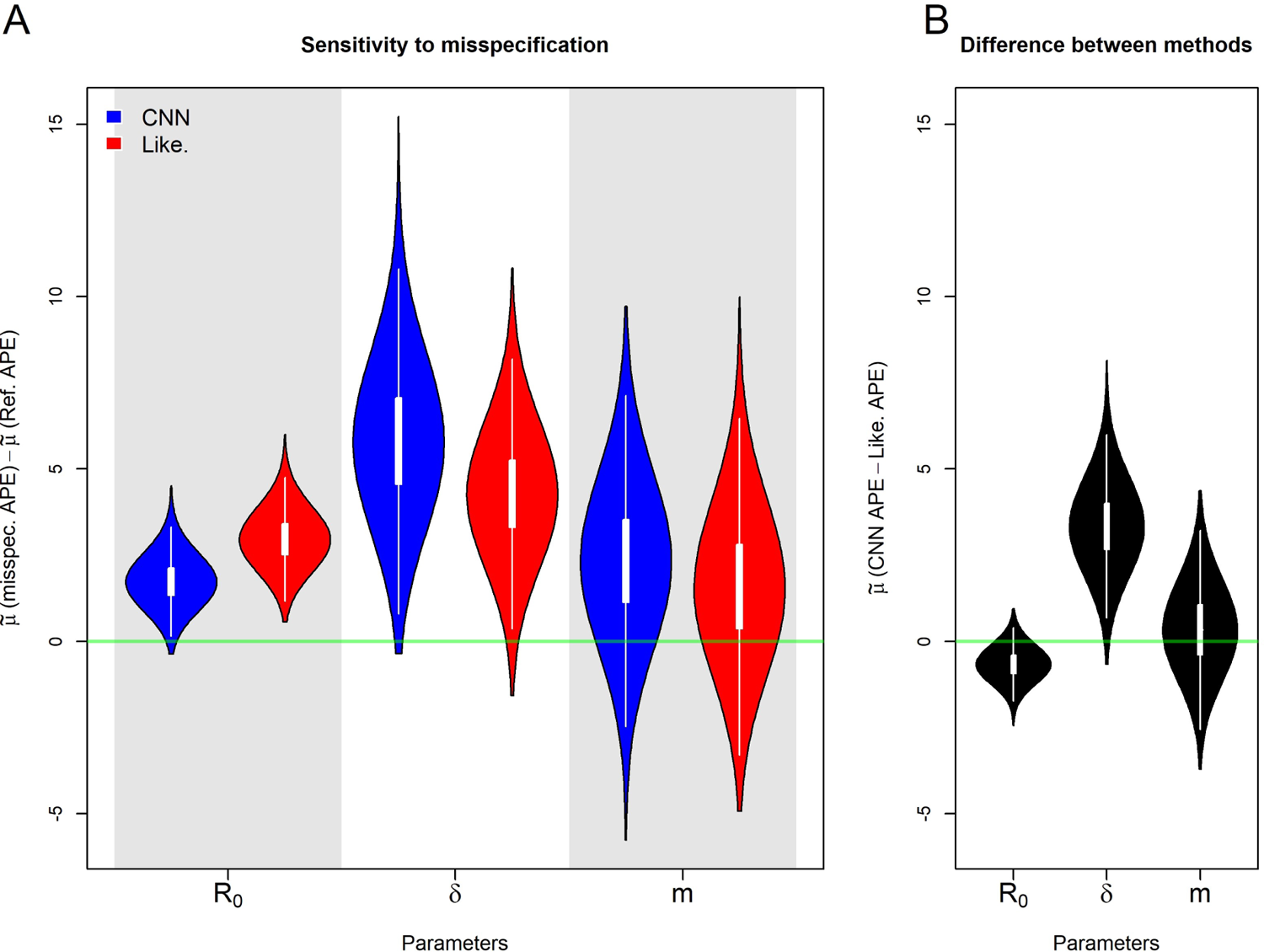
Posterior distributions of the median APE when the phylogenetic tree is incorrect. Details are the same as in S5

**Figure S14:**
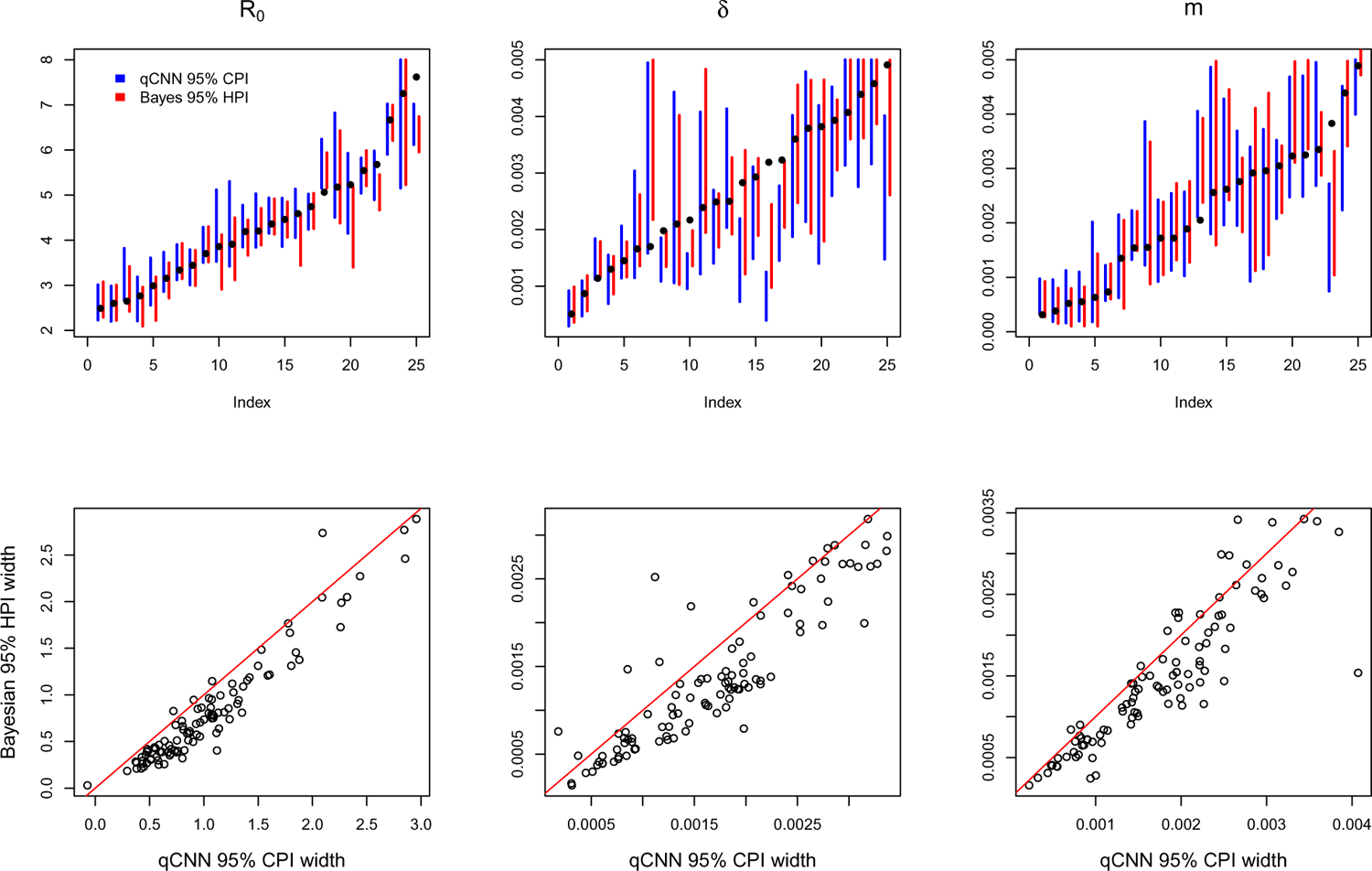
Comparison of CPI and HPI intervals for phylogeny error experiment. See SI Figure S4 for general details about plot.

